# Robust differentiation of human pluripotent stem cells into endothelial cells via temporal modulation of ETV2 with modified mRNA

**DOI:** 10.1101/2020.03.02.973289

**Authors:** Kai Wang, Ruei-Zeng Lin, Xuechong Hong, Alex H. Ng, Chin Nien Lee, Joseph Neumeyer, Gang Wang, Xi Wang, Minglin Ma, William T. Pu, George M. Church, Juan M. Melero-Martin

## Abstract

Human induced pluripotent stem cell (h-iPSC)–derived endothelial cells (h-iECs) have become a valuable tool in regenerative medicine. However, current differentiation protocols remain inefficient and lack reliability. Here, we describe a method for rapid, consistent, and highly efficient generation of h-iECs. The protocol entails the delivery of modified mRNA encoding the transcription factor *ETV2* at the intermediate mesodermal stage of differentiation. This approach reproducibly differentiated thirteen diverse h-iPSC lines into h-iECs with exceedingly high efficiency. In contrast, standard differentiation methods that relied on endogenous *ETV2* were inefficient and notably inconsistent. Our h-iECs were functionally competent in many respects, including the ability to form perfused vascular networks *in vivo*. Importantly, timely activation of *ETV2* was critical, and bypassing the mesodermal stage produced putative h-iECs with reduced expansion potential and inability to form functional vessels. Our protocol has broad applications and could reliably provide an unlimited number of h-iECs for vascular therapies.

## INTRODUCTION

Endothelial cells (ECs) are implicated in the pathogenesis of numerous diseases particularly due to their ability to modulate the activity of various stem cells during tissue homeostasis and regeneration (*1, 2*). Consequently, deriving competent ECs is central to many efforts in regenerative medicine. The advent of human induced pluripotent stem cells (h-iPSCs) created an exciting and non-invasive opportunity to obtain patient-specific ECs. However, differentiating h-iPSCs into ECs (herein referred to as h-iECs) with high efficiency, consistently, and in high abundance remains a challenge (*3*).

Current differentiation protocols are inspired by vascular development and rely on sequentially transitioning h-iPSCs through two distinct stages (referred to as stages 1 and 2 or S1-S2) (*4*). During S1, h-iPSCs differentiate into intermediate mesodermal progenitor cells (h-MPCs), a process regulated by Wnt and Nodal signaling pathways. In S2, h-MPCs acquire endothelial specification principally via VEGF signaling (*4*). Existing protocols, however, are far from optimal. Limitations stem from the inherent complexity associated with developmental processes. First, directing h-MPCs to solely differentiate into h-iECs is not trivial. Indeed, recent reports estimate that with the canonical S1-S2 approach, less than 10% of the differentiated cells may actually be *bona fide* h-iECs (*3*). In addition, achieving consistent differentiation in different h-iPSC lines continues to be a challenge (*5*). This dependency on cellular origin makes the clinical translation of h-iECs problematic.

The transcription factor E26 transformation-specific (ETS) variant 2 (*ETV2*) plays a non-redundant and indispensable role in vascular cell development (*6–9*). Importantly, expression of ETV2 is only required transiently. Recent studies have proposed reprograming somatic cells using transducible vectors encoding *ETV2* (*10–13*). Nevertheless, the efficiency of direct reprogramming somatic cells into ECs remains exceedingly low and achieving proper EC maturation requires long periods of time in culture. Alternatively, a few studies have recently proposed inducing *ETV2* expression directly on h-iPSCs to induce EC differentiation (*14–16*). However, to date, methods have relied on early activation of *ETV2* in the h-iPSCs, thus bypassing transition through an intermediate mesodermal stage. Also, the functional competence of the resulting h-iECs remains somewhat unclear.

Here, we sought to develop a protocol that enables more consistent and highly efficient differentiation of human h-iPSCs into h-iECs. We identified that a critical source of inconsistency resides in the inefficient activation of ETV2 during S2. To circumvent this constraint, we made use of chemically modified mRNA (modRNA), a technology that in recent years has improved the stability of synthetic RNA allowing its transfer into cells (and subsequent protein expression) *in vitro* and *in vivo* (*17*). We developed a synthetic modRNA to uniformly activate *ETV2* expression in h-MPCs, independently of VEGF signaling. As a result, conversion of h-MPCs into h-iECs occurred rapidly and robustly. Indeed, we reproducibly differentiated 13 different human h-iPSC clonal lines into h-iECs with high efficiency (>90%). Moreover, we demonstrated that these h-iECs were phenotypically and functionally competent in many respects, including their ability to form perfused vascular networks *in vivo*.

## RESULTS

### Rapid and highly efficient differentiation of human h-iPSCs into h-iECs

We developed a two-dimensional, feeder-free, and chemically defined protocol that relies on a timely transition of h-iPSCs through two distinct stages, each lasting 48 h. First is the conversion of h-iPSCs into h-MPCs. This step is similar to that in the standard S1-S2 differentiation protocol and thus is mediated by the activation of Wnt and Nodal signaling pathways using the glycogen synthase kinase 3 (GSK-3) inhibitor CHIR99021 (Fig. 1A). Second, we converted the h-MPCs into h-iECs. This step is substantially different than the S1-S2 protocol which relies on activation of endogenous *ETV2* via VEGF signaling. In contrast, our protocol used chemically modified RNA (modRNA) to deliver exogenous *ETV2* to h-MPCs via either electroporation or lipofection (Fig. 1A).

**Figure 1.**
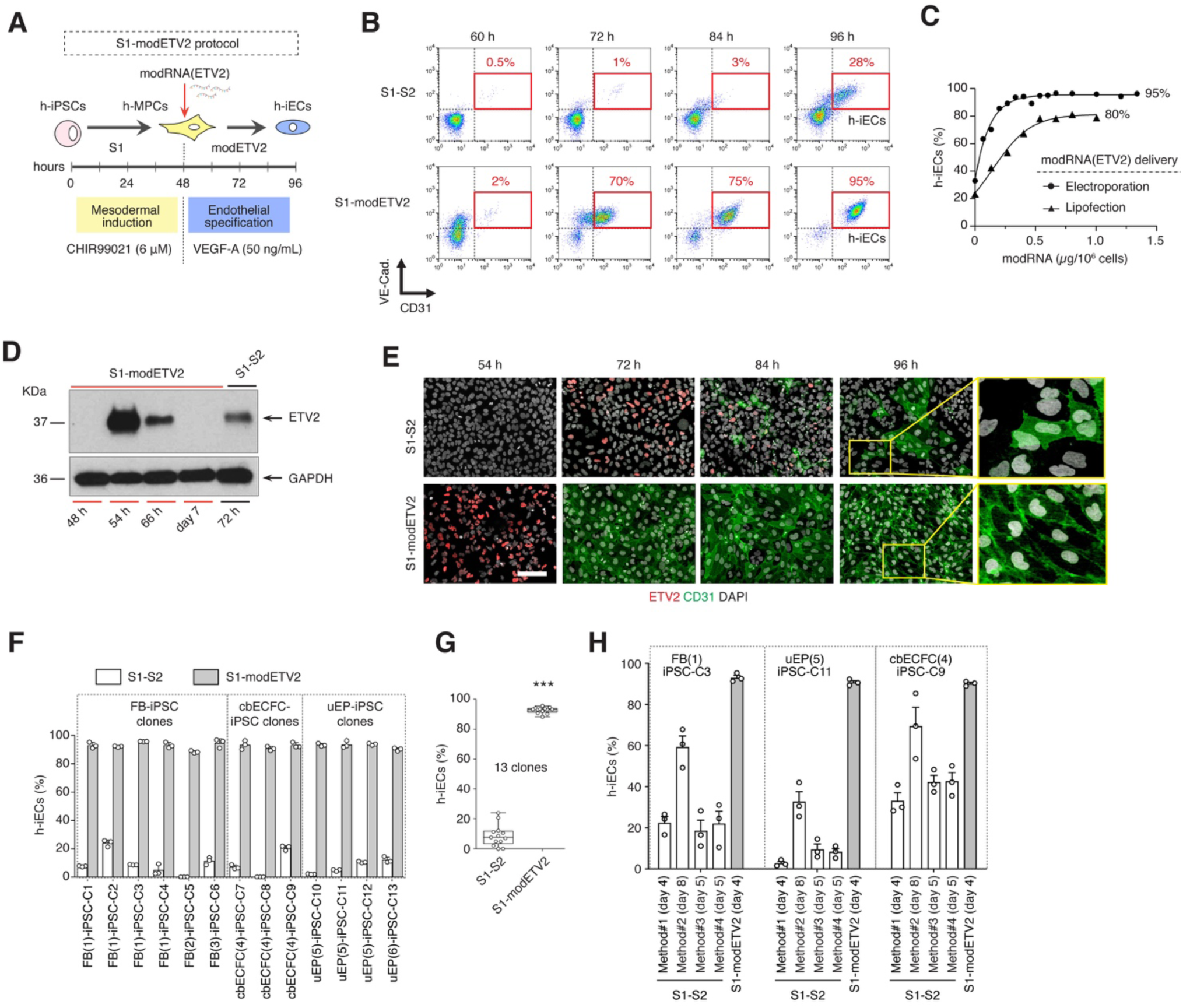
Robust endothelial differentiation of h-iPSCs. (**A**) Schematic of optimized two-stage endothelial differentiation protocol. Stage 1: conversion of h-iPSCs into h-MPCs mediated by the GSK-3 inhibitor CHIR99021. Stage 2: transfection of h-MPCs with modRNA encoding *ETV2* and culture in chemically defined medium. (**B**) Conversion efficiency of h-iPSCs into VE-Cadherin+/CD31+ h-iECs by flow cytometry. Time course comparison of the standard S1-S2 and the optimized S1-modETV2 protocols. (**C**) Effect of modRNA concentration on h-iPSC-to-h-iEC conversion efficiency at 96 h. Titration analysis of CD31+ cells by flow cytometry for both electroporation- and lipofection-based delivery of modRNA. (D) Western Blot analysis of ETV2 expression in differentiation. Lanes #1-4 correspond to cells at different time point of the S1-modETV2 protocol. Lane #5 corresponds to cells at day 3 (S2-72 h) of the standard S1-S2 protocol. (**E**) Time course immunofluorescence staining for ETV2 and CD31 in S1-S2 and S1-modETV2 protocols. Nuclei stained by DAPI. Scale bar, 100 μm. (**F**) Flow cytometry analysis of differentiation efficiency at 96 h in 13 h-iPSC clones generated from dermal fibroblasts (FB), umbilical cord blood-derived ECFCs (cbECFC), and urine-derived epithelial cells (uEP). (**G**) Differences in differentiation efficiency between S1-S2 and S1-modETV2 protocols for all 13 h-iPSC clones. Data correspond to percentage of CD31+ cells by flow cytometry. (**H**) Differences in differentiation efficiency between four alternative S1-S2 methodologies and the S1-modETV2 protocol for 3 independent h-iPSC clones. Bars represent mean ± s.d.; \*\*\**P* < 0.001.

Our customized two-step protocol (herein referred to as S1-modETV2) rapidly and uniformly converted human h-MPCs into h-iECs. Indeed, 48 h after transfection of h-MPCs with modRNA(*ETV2*), ∼95% of the cells were endothelial (VE-Cadherin+/CD31+ cells; Fig. 1B) (see fig. S1, A and B for controls accounting for electroporation with no modRNA and with modRNA(GFP)). In contrast, conversion during the standard S2 step (no modRNA) was slower and significantly less efficient, with less than 30% VE-Cadherin+/CD31+ cells at the same time point (Fig. 1B) Conversion efficiency was dependent on the amount of modRNA(ETV2) used. Titration analysis revealed that above 0.5 µg of modRNA(ETV2) per 10^6^ h-iPSCs, the percentage of h-iECs at 96 h was consistently ∼95% (using electroporation) and ∼75% (lipofection) (Fig. 1C; fig. S1, C and D).

Transfection with modRNA(ETV2) enabled rapid, transient, and uniform expression of ETV2, in contrast to delayed and sparse expression with the S1-S2 method (Fig. 1D-E). Broad expression of ETV2, in turn, resulted in uniform CD31 expression by 96 h (Fig. 1E). During the S1-S2 protocol, the presence of non-endothelial VE-Cadherin-/SM22+ cells was prominent at 96 h (fig. S1E). However, the occurrence of VE-Cadherin-/SM22+ cells was significantly reduced in our S1-modETV2 protocol (<3%), suggesting a more effective avoidance of alternative non-endothelial differentiation pathways (fig. S1E).

### Differentiation reproducibility with clonal h-iPSC lines from various cellular origins

Current S1-S2 differentiation protocols lack consistency between different h-iPSC lines. To address this limitation, we generated multiple human clonal h-iPSC lines from three distinct cellular origins corresponding to subcutaneous dermal fibroblasts (FB), umbilical cord blood-derived endothelial colony-forming cells (cbECFC), and urine-derived epithelial cells (uEP) (fig. S2A). All h-iPSCs were generated with a non-integrating episomal approach and validated by expression of pluripotent transcription factors OCT4, NANOG, and SOX2; and capacity to form teratomas in immunodeficient mice (fig. S2, B and C).

We generated 13 clones (referred to as C1-C13) to collectively represent variations due to different individual donors, cellular origins, and clone selection (fig. S2). All clones were subjected to both S1-S2 and S1-modETV2 differentiation protocols. As expected, the S1-S2 protocol produced a wide variation in efficiency, with h-iECs ranging from <1% to ∼24% (Fig. 1G-H). In addition, there were noticeable inconsistencies between clones from similar cellular origins but different donors (e.g., C2 vs. C5; and C10 vs. C13) and even between genetically-identical clones derived from the same h-iPSC line (e.g., C2 vs. C4; C8 vs. C9; and C10 vs. C12) (Fig. 1G). In contrast, differentiation under the S1-modETV2 protocol produced significantly higher efficiencies and eliminated inconsistencies between clones. Indeed, in all 13 clones, the percentage of CD31+ h-iECs at 96 h ranged between 88-96% irrespective of the donor and cellular origin from which the h-iPSC clones were derived, and there were no statistical differences in efficiency (Fig. 1G-H). Thus, the S1-modETV2 method was highly consistent in terms of differentiation efficiency across different clones.

To further corroborate these findings, we examined three additional S1-S2 methodologies corresponding to protocols described by Harding *et al*. 2017 (*18*) (Method#2; duration: 8 days), Sahara *et al*. 2014 (*19*) (Method#3; 5 days), and Patsch *et al*. 2015 (*4*) (Method#4; 5 days). These methods were compared to our S1-S2 method (referred to as Method#1; 4 days; Fig.1H) and the S1-modETV2 method. For this comparison, we used three independent h-iPSC clones corresponding to three different cellular origins: dermal fibroblasts (clone FB(1)-iPSC-C3); umbilical cord blood-derived endothelial colony-forming cells (cbECFC(4)-iPSC-C9), and urine-derived epithelial cells (uEP(5)-iPSC-C11) (Fig.1H). Examination of the efficiency revealed that all four S1-S2 differentiation protocols produced significantly lower efficiencies than the S1-modETV2 method. Moreover, depending on the h-iPSC clone used, there was a wide variation in efficiency among the four S1-S2 methods, with h-iECs ranging from ∼3% to ∼70% (Fig.1H). In contrast, differentiation under the S1-modETV2 protocol produced significantly higher efficiencies (89-95%) and eliminated inconsistencies between clones.

### Inefficient activation of endogenous *ETV2* in intermediate h-MPCs

To further evaluate the issue of inefficiency, we carried out a transcriptional examination of the standard S1-S2 differentiation protocol. As expected, conversion of h-iPSCs into h-MPCs coincided with transient activation of mesodermal transcription factors *MIXL1* and *TBXT* (Fig. 2A; fig. S3A). Likewise, differentiation of h-MPCs into h-iECs involved activation of *ETV2* (transiently) and then *ERG* (Fig. 2A; fig. S3A), consistent with previous vascular developmental descriptions (*6, 20*). However, there were significant differences when comparing the efficiencies of transcription factor activation. On one hand, activation of *TBXT* (which encodes for Brachyury) was robust and highly uniform at 48 h (∼97% Brachyury+ cells; Fig. 2B), suggesting that the conversion of h-iPSCs into h-MPCs is unlikely to account for the large inefficiency observed in the S1-S2 protocol. On the other hand, *ETV2* activation by 72 h was limited and far from uniform (∼33% ETV2+ cells; Fig. 2C), indicating inefficient conversion of h-MPCs into h-iECs. Control h-iPSCs in which *ETV2* was genetically abrogated using CRISPR-Cas9 (h-iPSC-ETV2^-/-^) displayed unaltered *TBX*T activation and mesodermal conversion, but were unable to activate *ETV2* and, in turn, incapable of initiating S2 (Fig. 2B-C; fig. S4, C and D). Because *ETV2* expression is governed by VEGF signaling (*21*), we examined whether increasing the concentration of supplemented VEGF could improve its inefficient activation during S1-S2. However, we found that beyond 50 ng/mL VEGF failed to further increase the proportion of ETV2+ cells and thus the subsequent conversion to CD31+ h-iECs (Fig. 2D; fig. S4, F and G).

**Figure 2.**
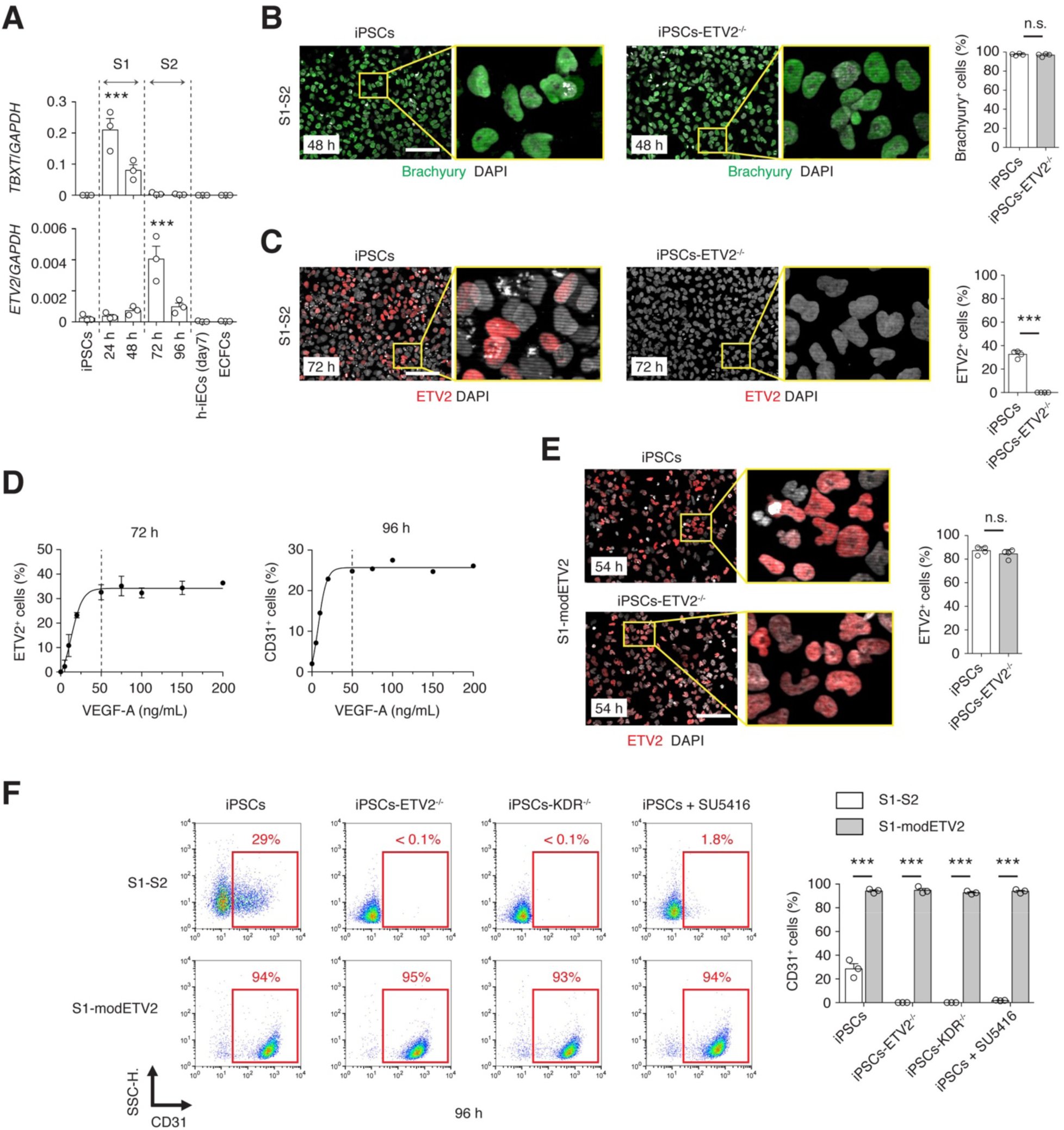
Inefficient activation of endogenous *ETV2* in intermediate h-MPCs during the standard S1-S2 differentiation protocol. (**A**) Time course analysis of mRNA expression (qRT– PCR) of transcription factors *TBXT* (mesodermal commitment) and *ETV2* (endothelial commitment) during the standard S1-S2 differentiation protocol. Relative fold change normalized to *GAPDH* expression. (**B**) Immunofluorescence staining for Brachyury in h-iPSCs at 48 h during the S1-S2 protocol. h-iPSCs lacking endogenous *ETV2* (h-iPSCs-ETV2^-/-^) served as control. Nuclei stained by DAPI. Scale bar, 100 μm. Percentage of Brachyury+ cells at day 1. (**C**) Immunofluorescence staining for ETV2 in h-iPSCs at 72 h during the S1-S2 protocol. h-iPSCs-ETV2^-/-^ served as control. Nuclei stained by DAPI. Scale bar, 100 μm. Percentage of ETV2+ cells at day 3. (**D**) Effect of VEGF-A concentration on the percentages of ETV2+ cells at 72 h and CD31+ cells at 96 h during the S1-S2 protocol measured by immunofluorescence staining (ETV2) and flow cytometry (CD31). (**E**) Immunofluorescence staining for ETV2 in h-iPSCs during the optimized S1-modETV2 protocol. h-iPSCs-ETV2^-/-^ served as control. Nuclei stained by DAPI. Scale bar, 100 μm. Percentage of ETV2+ cells after transfection with modRNA. (**F**) Conversion efficiency of h-iPSCs into CD31+ h-iECs by flow cytometry. Comparison of the standard S1-S2 and the S1-modETV2 protocols. h-iPSCs-ETV2^-/-^, h-iPSCs-KDR^-/-^, and h-iPSCs treated with the VEGFR2 inhibitor SU5416 served as controls. In panels b, c, and e, bars represent mean ± s.d.; n = 4; n.s. = no statistical differences and \*\*\**P* < 0.001 between h-iPSCs and h-iPSCs-ETV2^-/-^. In panel f, n = 3; \*\*\**P* < 0.001 between indicated groups.

Collectively, we found that while conversion of h-iPSCs into *TBXT*+ h-MPCs occurs very efficiently (>95%), activation of endogenous *ETV2* in h-MPCs is clearly limited (∼30%) during the S1-S2 protocol and did not improve by simply increasing VEGF concentration. Thus, we concluded that in order to improve the conversion of h-iPSCs into h-iECs, emphasis should be put on finding new means to effectively activate *ETV2* in the intermediate h-MPCs.

### Transient expression of exogenous *ETV2* uniformly convert h-MPCs into h-iECs

Our approach to more uniformly activate *ETV2* in h-MPCs is to use modRNA. Indeed, 6 h after transfection of h-MPCs with modRNA(ETV2), >85% of the cells expressed ETV2 (Fig. 2E), a significant increase from the mere ∼30% observed in the S1-S2 protocol. Of note, h-iPSC-ETV2^-/-^ also displayed widespread ETV2 expression after transfection, indicating that activation was independent of endogenous *ETV2* (Fig. 2E). This robust expression of *ETV2* produced high rates of endothelial specification and efficient conversion into h-iECs in both unmodified h-iPSCs and h-iPSC-ETV2^-/-^ with the S1-modETV2 differentiation protocol (Fig. 2F). In contrast, with the S1-S2 protocol, the differentiation process was less efficient and completely dependent on endogenous *ETV2* expression (h-iPSC-ETV2^-/-^ failed to produce h-iECs) (Fig. 2F; fig. S4E). Moreover, the S1-modETV2 protocol maintains the differentiation process independent of VEGFR-2 signaling. Indeed, h-iPSCs in which *KDR* (which encodes for VEGFR-2) was genetically abrogated using CRISPR-Cas9 (h-iPSC-KDR^-/-^) displayed an unaltered ability to differentiate into h-iECs (93% at 96 h) with the S1-modETV2 method but were incapable of differentiating (<0.1%) with the standard S1-S2 protocol (Fig. 2F; fig. S4, A and B). Likewise, chemical abrogation of VEGFR2 signaling with the inhibitor SU5416 impaired the differentiation of h-iPSCs into h-iECs with the S1-S2 protocol but not with the S1-modETV2 protocol (<2% and 94% h-iECs at 96 h, respectively) (Fig. 2F).

Taken together, we showed that delivery of modRNA(ETV2) is an effective means to robustly and transiently express ETV2 in intermediate h-MPCs, which in turn initiates widespread conversion into h-iECs. Our S1-modETV2 protocol renders the differentiation process independent of VEGFR-2 signaling and of endogenous ETV2, thus overcoming one of the main limitations in current protocols.

### Time of ETV2 activation affects the transcriptional profile of h-iECs

Previous studies have suggested that inducing *ETV2* expression directly on h-iPSCs could generate h-iECs without transition through an intermediate mesodermal stage (*14*). However, it remains unclear whether this strategy produces functionally competent h-iECs. To address this question, we generated putative h-iECs by transfecting h-iPSCs with modRNA(ETV2) (protocol herein referred to as early modETV2) (Fig. 3A). This method converted human h-iPSCs into CD31+ cells rapidly and efficiently (Fig. 3B, fig. S5), which is consistent with previous reports (*16*). Moreover, conversion was dependent on the concentration of modRNA(ETV2) and reproducible in all h-iPSCs clones tested, irrespective of the donor and cellular origin of the clones (fig. S5, C and D). Transfection of h-iPSCs with modRNA(ETV2) enabled early and transient expression of ETV2 (fig. S5, E and F), which occurred without previous significant expression of TBXT, suggesting a bypass of the intermediate mesodermal stage (fig. S6). With this approach, there was a remnant of undifferentiated (non-transfected) CD31-/OCT4+ cells at 48 h, which was deemed undesirable (fig. S5E). Nonetheless, repeated subculture and purification of CD31+ cells largely mitigated this concern.

**Figure 3.**
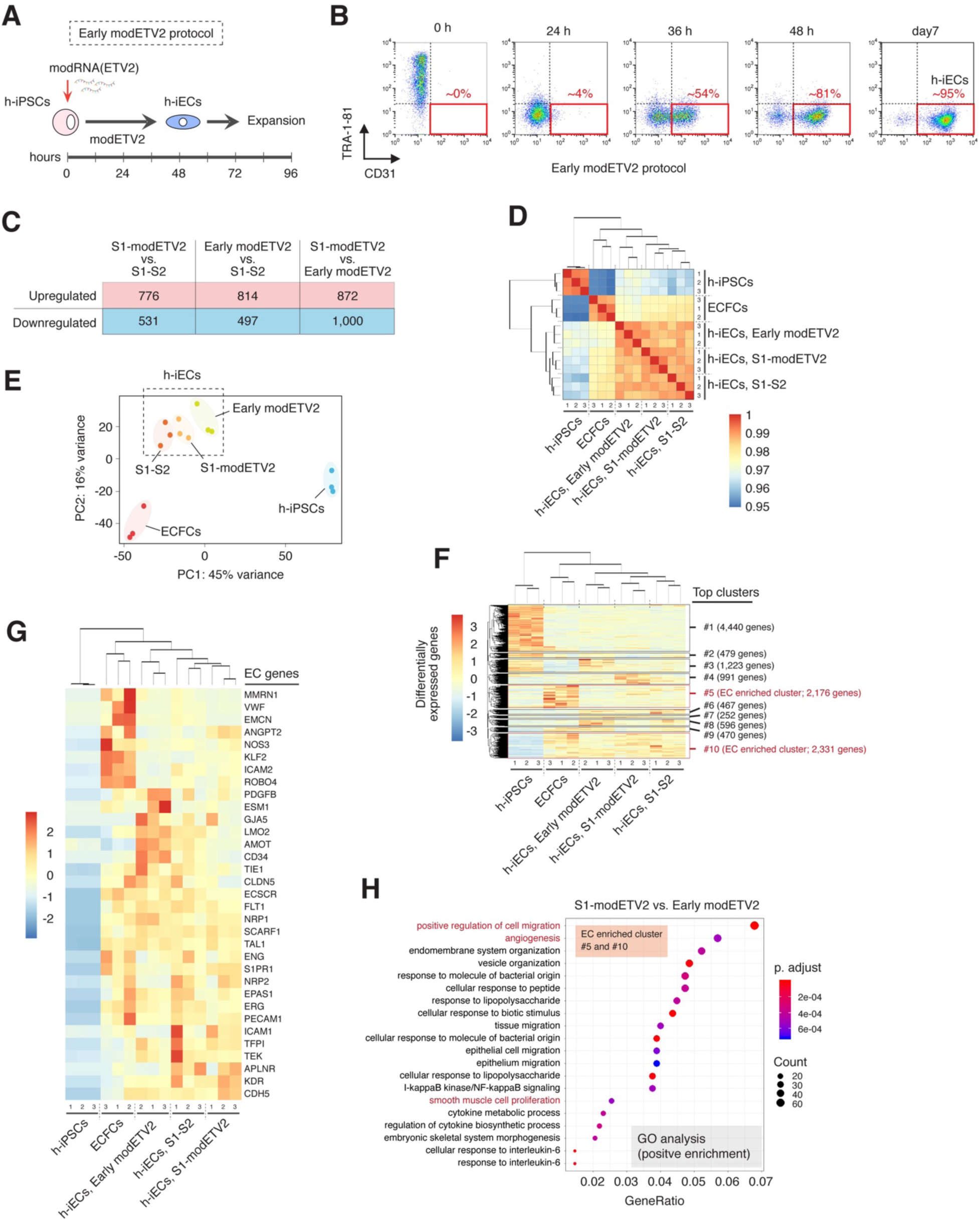
Transcriptional analysis of h-iECs obtained from various differentiation protocols. (**A**) Schematic of protocol for early transfection of h-iPSCs with modRNA encoding *ETV2*. (**B**) Conversion efficiency h-iPSCs into CD31+ cells by flow cytometry using the early modETV2 protocol. (**C-H**) RNAseq analysis across multiple h-iECs samples generated from three independent h-iPSC lines using all three differentiation protocols. Human ECFCs and the parental undifferentiated h-iPSCs served as positive and negative controls, respectively. (**C**) Number of differentially expressed genes between h-iECs samples from each differentiation protocol. (**D**) Pairwise correlation based on Pearson coefficients between all samples. (**E**) Principal component analysis. (**F**) Heatmap and hierarchical clustering analysis of global differentially expressed genes. (**G**) Heatmap and hierarchical clustering analysis of selected EC-specific genes. (**H**) GO analysis between h-iECs generated with the S1-modETV2 and the early modETV2 differentiation protocols. Analysis carried out with differentially expressed genes from EC clusters #5 and #10 based on (f). Genes displayed correspond to positive enrichment for h-iECs generated with the S1-modETV2 protocol.

To further elucidate potential differences between h-iECs generated from our S1-modETV2, the S1-S2, and the early modETV2 protocols, we performed RNAseq analysis across multiple h-iECs samples generated from three independent h-iPSC lines using all three differentiation protocols. Human ECFCs and the parental undifferentiated h-iPSCs served as positive and negative controls, respectively. Globally, there were thousands of differentially expressed genes across all the h-iEC groups (Fig. 3C; fig. S7A). Nevertheless, hierarchical clustering analysis of differentially expressed genes revealed 1) proximity between all the h-iEC groups, and 2) that h-iECs were transcriptionally closer to ECFCs than to h-iPSCs (Fig. 3F). These patterns of hierarchical association were confirmed by pairwise correlation (Fig. 3D) and principal component analyses (Fig. 3E). Moreover, analysis of selected endothelial and pluripotent genes confirmed that all groups of h-iECs were transcriptional more consistent with an endothelial phenotype than with the parental pluripotent state (Fig. 3G; fig. S7B) (Please note that Fig. 3G is not intended as a comprehensive list of EC genes). Importantly, our analysis also revealed that among h-iECs, there was more transcriptional proximity between h-iECs generated from the standard S1-S2 protocol and our S1-modETV2 method (Pearson’s correlation coefficient r = 0.987) than between h-iECs derived from the early modETV2 protocol and the other h-iEC groups (Fig. 3D).

To gain more insight into the transcriptional differences, we carried out gene ontology (GO) enrichment analysis between h-iECs generated with our S1-modETV2 and the early modETV2 differentiation protocols. Of note, analysis of all differentially expressed genes revealed that h-iECs generated with our S1-modETV2 displayed significant enrichment in genes associated with positive regulation of cell migration (fig. S7C). Moreover, a GO analysis was performed with differentially expressed genes from EC clusters #5 and #10 (cluster elucidated by hierarchical clustering analysis; Fig. 3F). Results indicated positive enrichment in h-iECs generated with our S1-modETV2 of genes associated with not only cell migration but also angiogenesis and smooth muscle proliferation (Fig. 3H; fig. S7D), suggesting differences in genes affecting critical vascular function.

### Early activation of ETV2 renders putative h-iECs with impaired functionality

Next, we examined whether the transcriptional differences observed between h-iECs generated from different protocols affected their capacity to function as proper ECs. Specifically, we compared h-iECs that were generated with the standard S1-S2, our S1-modETV2, and the early modETV2 protocols. Of note, ETV2 expression is transient in both differentiation protocols, and thus at the time of h-iEC characterization, ETV2 expression was completely absent (Fig. 1, D and E). Human cord blood-derived ECFCs served as control for *bona fide* ECs. First, we assessed the capacity to grow in culture. Previous studies have shown mixed results with regards to the expansion potential of h-iECs and currently there is no consensus on this issue (*22*). We observed that h-iECs generated with our S1-modETV2 protocol were easily expanded in culture for a period of 3 weeks, with an average expansion yield of ∼70-fold (Fig. 4A). This yield was significantly higher than that of h-iECs produced with the S1-S2 differentiation protocol (∼20-fold), which was mainly attributed to differences in efficiency during the initial 4 days of differentiation (Fig. 4A). More striking, however, was the lack of expansion displayed by h-iECs generated with the early modETV2 protocol. Notwithstanding the high differentiation efficiency of this method (Fig. 3A), these putative h-iECs ceased proliferating after approximately two weeks in culture with only a modest overall yield of ∼2-fold (Fig. 4A). In addition, it is important to note that h-iECs obtained by the S1-modETV2 method retained an endothelial phenotype, with typical cobblestone-like morphology along their expansion in culture, did not express the pluripotent marker *OCT4,* expressed numerous EC markers at the mRNA and protein levels, and showed affinity for the binding of *Ulex europaeus agglutinin I* (UEA-I) lectin (fig. S8). Indeed, although it is certainly possible that further maturation occurs during expansion *in vitro*, h-iECs fundamentally remained endothelial when one considers a plethora of EC markers, even though the level of expression of some individual genes (such as *NOS3* and *vWF*) varied over time (fig. S8C). Examination at days 4, 11, and 21 during expansion revealed that h-iECs remained fairly pure (>95% VE-cadherin+/CD31+ cells), maintained expression of EC markers at the mRNA and protein levels, and remained negative for *POU5F1* (OCT4) and α-Smooth muscle actin (α-SMA) (fig. S8D).

**Figure 4.**
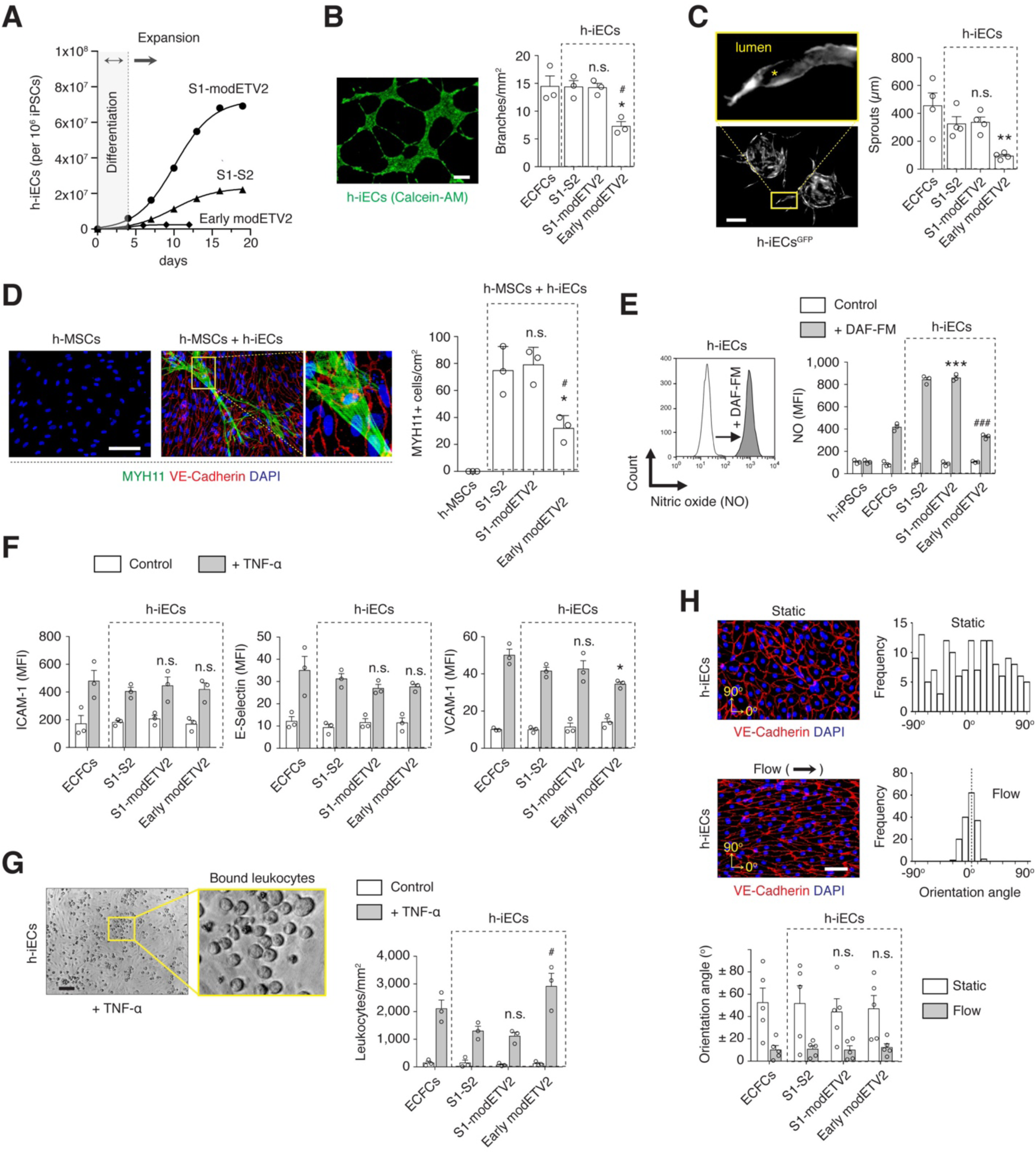
Functional properties of h-iECs. (**A**) Expansion potential of h-iECs derived by the standard S1-S2, optimized S1-modETV2, and early modETV2 protocols. Cumulative cell number measured over time in serially passaged h-iECs, starting with 10^6^ h-iPSCs. All h-iECs were purified as CD31+ cells at day 4. (**B**) Capillary-like networks formed by h-iECs on Matrigel. Live cells stained by Calcein-AM. Scale bar, 200 μm. The ability to form capillary-like networks was quantified and expressed as total number of branches per mm^2^. (**C**) Sprouts formation by spheroids formed with h-iECs^GFP^ and h-MSCs and embedded in fibrin gel for 4 days. The ability to form lumenal sprouts was quantified and expressed as total length per field. Scale bar, 200 μm. (**D**) Induction of smooth muscle cell differentiation of h-MSCs by h-iECs. Representative immunofluorescent images of h-MSCs in the absence or presence of h-iECs. Differentiation was assessed by the expression of smooth muscle myosin heavy chain 11 (MYH11). Expression of VE-Cadherin was used to detect h-iECs and DAPI for cell nuclei. Scale bar, 100 μm. Quantification of smooth muscle myogenic differentiation as number of h-MSCs expressing MYH11 per unit of culture area. (**E**) Nitric oxide (NO) production by h-iECs detected by flow cytometry as mean fluorescence intensity upon exposure to 4-amino-5-methylamino-2’,7’-difluorofluorescein diacetate (DAF-FM). Cells without exposure to DAF-FM served as negative control. (**F**) Upregulation of leukocyte adhesion molecules ICAM-1, E-Selectin, and VCAM-1 in h-ECs measured as mean fluorescence intensity by flow cytometry upon exposure to TNF-α. Cells not exposed to TNF-α served as negative control. (**G**) Representative brightfield images of h-iECs with an increased number of bound leukocytes after TNF-α treatment. Scale bar, 50 µm. Quantification of bound leukocytes per mm^2^ upon exposure to TNF-α. Cells without exposure to TNF-α served as negative control. (**H**) Capacity of h-iEC to align in the direction of flow. Representative immunofluorescent images of h-iECs under static and flow conditions. Cells stained by CD31 and nuclei by DAPI. Scale bar, 100 μm. Cell alignment quantified as frequency of cell orientation angle in histogram plots. 0° represents the direction of flow. In all panels, bars represent mean ± s.d.; n = 3; \**P* < 0.05, \*\**P* < 0.01, \*\*\**P* < 0.001 between h-iECs and ECFCs. ^#^*P* < 0.05, ^##^*P* < 0.01, ^###^*P* < 0.001 compared to h-iECs generated by S1-S2 protocol. n.s. = no statistical differences compared to both ECFCs and h-iECs generated by S1-S2 protocol.

We then evaluated the performance of h-iECs using an array of standard endothelial functional assays, including ability to: (i) assemble into capillary-like structures (Fig. 4B); (ii) launch angiogenic sprouts with proper lumens (Fig. 4C); (iii) induce smooth muscle differentiation of human mesenchymal stem cells (h-MSCs) (Fig. 4D); (iv) produce nitric oxide (NO) (Fig. 4E); (v) up-regulate expression of leukocyte adhesion molecules (E-selectin, ICAM-1, and VCAM-1) upon exposure to tumor necrosis factor-alpha (TNF-α) (Fig. 4F); (vi) up-regulate leukocyte binding upon exposure to TNF-α (Fig. 4G); and (vii) sense and adapt to shear flow, aligning to the direction of flow (Fig. 4H). Collectively, this comprehensive examination confirmed that h-iECs generated with both the S1-S2 and our S1-modETV2 differentiation protocols were functionally very similar, with no statistically significant differences between both groups in any of the assays (Fig 4, B to H). In addition, both h-iEC groups were comparable to the control ECFCs (the only exception was higher NO production by h-iECs; *P*<0.001; Fig. 4E), thus suggesting adequate endothelial function.

In contrast, h-iECs generated with the early modETV2 protocol showed signs of impaired functionality. When compared to the control ECFCs, these h-iECs appeared competent in some fundamental capacities such as the ability to regulate leukocyte adhesion upon an inflammatory stimulus, and the capacity to align in the direction of flow. However, h-iECs generated with the early modETV2 protocol displayed quantitative deficiencies in several important respects, including a significantly lower ability to assemble into capillary-like structures (*P*<0.05; Fig. 4B), to launch proper angiogenic sprouts (*P*<0.01; Fig. 4C), and to induce smooth muscle differentiation of h-MSCs (*P*<0.05; Fig. 4D). These differences were also statistically significant when compared to h-iECs generated with both the S1-S2 and our S1-modETV2 differentiation protocols, suggesting certain fundamental phenotypic differences between h-iECs generated from the different protocols.

### Timely activation of ETV2 is critical for proper vascular network-forming ability

Lastly, we examined the capacity of the different h-iECs to assemble into functional blood vessels *in vivo* (Fig. 5). To this end, we used our model of vascular network formation in which human ECs are combined with supporting MSCs in a hydrogel, and the grafts are then implanted into immunodeficient NOD-SCID mice (*23*). After 7 days *in vivo*, macroscopic examination of the explants suggested differences in the degree of vascularization between implants containing different types of h-iECs (*n* = 5) (Fig. 5A). Histological (H&E) analysis revealed that grafts with h-iECs generated with the S1-modETV2 protocol had an extensive network of perfused microvessels (Fig. 5B, left). These microvessels were primarily lined by the h-iECs, as confirmed by the expression of human-specific CD31 (Fig. 5E, left) and mouse-specific CD31 (Fig. 5G), by the affinity for UEA-I (Fig. 5D, left), and by the use of gfp-labeled h-iECs (fig. S9D). Moreover, these human lumens contained mouse erythrocytes (Fig. 5B, left), indicating formation of functional anastomoses with the host circulatory system. The presence of perfusion was further assessed by the infusion of biotinylated UEA-I, which specifically bounded to the lumen of the human blood vessels, confirming that they were connected to the mouse circulation (Fig. 5H). In contrast, the number of perfused human vessels in grafts with h-iECs generated with the early modETV2 protocol was exceedingly low (Fig. 5B, right). These h-iECs remained organized as lumenal structures (Fig. 5, D and E, right), but the lumens were rarely perfused (Fig. 5B, right). Indeed, microvessel density in grafts containing h-iECs from the early modETV2 protocol was significantly lower than in any other group (Fig. 5C). Of note, there were no significant differences in vessel density between grafts formed with h-iECs from the S1-modETV2 protocol, the S1-S2 protocol (both of which underwent transition through mesodermal intermediates) and the control ECFCs.

**Figure 5.**
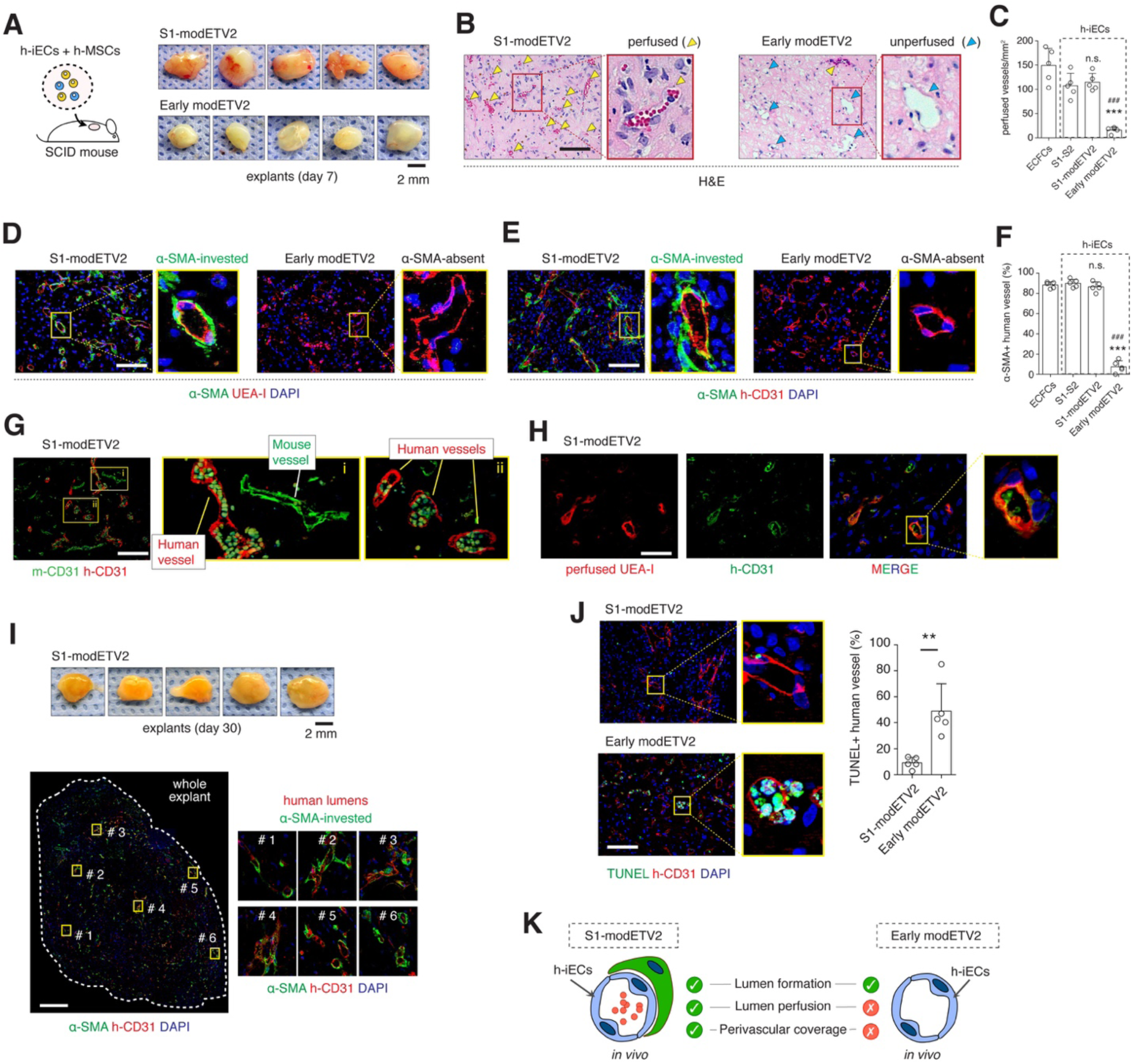
In vivo vascular network-forming ability of h-iECs. (**A**) Schematic of microvascular graft models. Grafts were prepared by combining h-iECs with h-MSCs in hydrogels and were then subcutaneously implanted into SCID mice for 7 days. Grafts contained h-iECs that were generated by either the S1-modETV2 or the early modETV2 protocol. Images are macroscopic views of the explanted grafts at day 7. (**B**) Haematoxylin and eosin (H&E) staining of explanted grafts after 7 days *in vivo*. Perfused vessels were identified as luminal structures containing red blood cells (RBCs) (yellow arrowheads). Blue arrowheads represent unperfused luminal structures. (**C**) Density of perfused blood vessels on day 7. Groups include grafts with h-iECs that were generated by either the standard S1-S2, the optimized S1-modETV2, or the early modETV2 protocol. Grafts with ECFCs served as control. (**D-E**) Immunofluorescence staining of explanted grafts after 7 days *in vivo*. Human lumens stained by (**D**) UEA-I and (**E**) h-CD31. Perivascular coverage stained by α-SMA. Nuclei stained by DAPI. Scale bar, 100 μm. Grafts with h-iECs from the S1-modETV2 protocol had an extensive network of α-SMA-invested human lumens, whereas grafts with h-iECs from the early modETV2 protocol presented human lumens with no perivascular coverage. (**F**) Percentage of human lumens with α-SMA+ perivascular coverage in explanted grafts at day 7. Groups include grafts with h-iECs that were generated by either the standard S1-S2, the optimized S1-modETV2, or the early modETV2 protocol. Grafts with ECFCs served as control. (**G**) Distinguishing human and mouse vessels by immunofluorescence. Immunofluorescence staining of explanted grafts after 7 days *in vivo*. Mouse lumens stained by anti-mouse-CD31 antibody (m-CD31; green). Human lumens stained by anti-human-CD31 antibody (h-CD31; red). Please note that the red blood cells within the lumen had auto-fluorescence. Scale bar, 100 μm. (**H**) Perfusion through human vessels reveled by the binding of infused biotinylated UEA-I followed by staining with Texas Red-conjugated Streptavidin (perfused UEA-1; red). Human vessels also stained by anti-human-CD31 antibody (h-CD31; green). Scale bar, 50 μm. (**I**) Grafts with h-iECs generated by the S1-modETV2 protocol were explanted at day 30. Images are macroscopic views of the explanted grafts. Immunofluorescence staining of explanted grafts by h-CD31 and α-SMA. Nuclei stained by DAPI. Scale bar, 500 μm. (**J**) Immunofluorescence staining by TUNEL and h-CD31 of explanted grafts after 7 days *in vivo*. Nuclei stained by DAPI. Scale bar, 100 μm. Percentage of human lumens that were TUNEL+ in explanted grafts at day 7. Groups include grafts with h-iECs generated by the S1-modETV2 and the early modETV2 protocols. Bars represent mean ± s.d.; n = 5; \*\**P* < 0.01. (**K**) Schematic of in vivo vascular network-forming ability of h-iECs generated by either the optimized S1-modETV2 or the early modETV2 protocols. In **C** and **F**, bars represent mean ± s.d.; n = 5; \*\*\**P* < 0.001 between h-iECs and ECFCs. ^###^*P* < 0.001 compared to h-iECs generated by S1-S2 protocol. n.s. = no statistical differences compared to both ECFCs and h-iECs generated by S1-S2 protocol.

It is important to note that although co-transplantation with MSCs facilitates engraftment, h-iECs derived from the S1-modETV2 protocol were also able to engraft and form perfused vessels when implanted alone, without MSCs (fig. S9). Indeed, grafts containing h-iECs alone became vascularized in 7 days and histological analysis confirmed the presence of numerous microvessels lined by the h-iECs (fig. S9, A to C).

We also examined the presence of mural cell investment around the newly-formed human vessels, a hallmark of proper vessel maturation and stabilization (*24*). There was a striking difference between h-iECs generated with the S1-modETV2 protocol and those generated with the early modETV2 with regard to perivascular investment (Fig. 5D-F). In grafts with h-iECs from the S1-modETV2 protocol, the large majority (∼87%) of the human vessels had proper coverage by perivascular cells expressing α-smooth muscle actin (α-SMA) (Fig. 5, D and E, left; Fig. 5F). This high percentage of perivascular coverage is to be expected by day 7 in this model (*25*), and vessels formed by control ECFCs consistently displayed high coverage (Fig. 5F). In contrast, grafts containing h-iECs from the early modETV2 protocol had only ∼8% of their human vessels covered by α-SMA+ cells (Fig. 5, D and E, right; Fig. 5F). These h-iECs were able to engraft and self-assemble into recognizable lumenal structures, but these structures lacked perivascular cells indicating inadequate maturation. Moreover, TUNEL staining of explants at day 7 revealed signs of apoptosis in a large percentage of human vessels lined by h-iECs from the early modETV2 protocol (Fig. 5J), an indication of vessel instability. On the other hand, 30 days after implantation, grafts that used h-iECs generated with the S1-modETV2 protocol still contained extensive and uniform networks of human vessels with proper perivascular coverage (Fig. 5I).

Taken all together, we demonstrated that during the differentiation of h-iPSCs into h-iECs, the ETV2 activation stage is critical. With our optimized S1-modETV2 protocol, activation of ETV2 occurred at the intermediate mesodermal stage, which produced h-iECs that were phenotypically and functionally competent. However, bypassing transition through the mesodermal stage by early activation of ETV2 produced putative h-iECs with a transcriptional profile further away from that of *bona fide* ECs, and, more importantly, with impaired functionality.

## DISCUSSION

Here, we developed a novel protocol that enables highly efficient differentiation of human h-iPSCs into competent h-iECs. The protocol entails a total differentiation period of 4 days and comprises two steps: 1) differentiation of h-iPSCs into intermediate h-MPCs; and 2) conversion of h-MPCs into h-iECs upon delivery of modRNA encoding *ETV2*. We showed that this S1-modETV2 approach allows widespread expression of ETV2 throughout the entire h-MPC population, thus overcoming one of the main hurdles of current protocols. Using our customized protocol, we reproducibly and efficiently differentiated 13 different human h-iPSC clonal lines into h-iECs. In all cases, we produced h-iECs at exceedingly high purity irrespective of the h-iPSC donor and cellular origin, and there were no statistical differences in efficiency. Of note, this high efficiency and reproducibility were absent when we used the standard S1-S2 protocol, which relies on VEGF signaling for endogenous *ETV2* activation. In addition, we were able to expand the resulting h-iECs with ease, obtaining an average h-iEC-to-h-iPSC ratio of ∼70-fold after 3 weeks in culture. More importantly, we demonstrated that our h-iECs were phenotypically, transcriptionally, and functionally consistent with *bona fide* ECs, including a robust ability to form perfused vascular networks *in vivo*.

Over the last decade, refinements to the standard S1-S2 differentiation protocol have steadily improved efficiency. Improvements have included, for example, the inhibition of the Notch and the TGF-β signaling pathways, the activation of protein kinase A or the synergistic effects of VEGF and BMP4 during S2 (*18, 19, 26*). However, most of these advances have been largely incremental, and consensus holds that the differentiation of h-iPSCs into h-iECs remains somewhat inconsistent (*27*). The incorporation of BMP4 during S2 was shown to produce a significant improvement in differentiation efficiency; however, the mechanism behind this improvement remains unknown and thus it is unclear whether this approach can consistently produce high efficiency across multiple clonal iPSC lines, independently of their cellular origin (*18*). One of the major difficulties is related to the necessary transition through the intermediate h-MPCs, which serve as common progenitors to not only h-iECs but also to other end-stage mesodermal cell types (*28, 29*). Thus, directing h-MPCs to solely differentiate into h-iECs is a challenge. Indeed, a recent study that used single-cell RNA analysis revealed that after S1-S2, non-endothelial cell populations (including, cardiomyocytes and vascular smooth muscle cells) were in fact predominant among the differentiated cells, and less than 10% were actually identified as *bona fide* ECs (*3*). Studies have also shown that EC specification is dictated by a transient activation of *ETV2*, which in turn depends on VEGF signaling (*21, 30*). However, our study has revealed that the activation of endogenous *ETV2* during S2 is inherently inefficient and increasing the concentration of VEGF can only improve this constraint to a certain degree. This limited ability of exogenous VEGF to enhance efficiency could be explained by the fact that VEGF has also been shown to promote h-iPSC differentiation into other mesodermal fates, including cardiac progenitor cells, cardiomyocytes and hepatic-like cells (*3, 28, 31–33*). Thus, in order to improve efficiency, VEGF activation of *ETV2* must be accompanied by inhibition of all other competing fates, which is not trivial. Our approach, however, circumvents this challenge. By using modRNA, we were able to transiently express ETV2 in a high percentage of h-MPCs and independently of VEGF signaling. This, in turn, allowed widespread conversion into h-iECs thereby eliminating the problem of inefficiency. In addition, our study provides an important new insight: that timely activation of *ETV2* is critical, and that bypassing the intermediate mesodermal stage is detrimental. Indeed, h-iECs generated by our S1-modETV2 methodology displayed proper blood vessel-forming ability *in vivo*, whereas putative h-iECs generated by the early modETV2 approach displayed impaired functionality and were unable to robustly form perfused vessels with adequate perivascular stability (Fig. 5K).

Current protocols are also limited by inconsistent results among different h-iPSC lines. Indeed, a recent study examined genetically identical h-iPSC clonal lines that were derived from various tissues of the same donor and found that by following the standard S1-S2 protocol, both differentiation efficiency and gene expression of the resulting h-iECs varied significantly depending on the source of h-iPSCs (*5*). In our study, we have also found inconsistencies in differentiation efficiency between h-iPSC clonal lines with different cellular origins, including lines with identical genetic make-up and lines derived from the same tissues in different donors. This lack of consistency is certainly undesirable from a clinical translation standpoint (*22*). Also, dependency on cellular origin may explain why published results on differentiation efficiency are often mixed and rely on selecting h-iPSC clones that are particularly attuned to EC differentiation. Our method eliminates this uncertainty as it consistently produces high efficiency, irrespective of the donor and cellular origin from which the h-iPSC clones are derived.

In summary, we have developed a protocol that enables highly efficient and reliable differentiation of human h-iPSCs into competent h-iECs. The protocol is simple, rapid, and entails delivery of modified mRNA encoding the transcription factor *ETV2* at the intermediate mesodermal stage of differentiation. In terms of added benefits, our method provides two key advantages over current published protocols. First, it significantly increases reproducibility among different clones of h-iPSCs and thus eliminates the need for selecting clones that are particularly attuned to activation of endogenous *ETV2* during differentiation. Second, our method yields exceedingly high differentiation efficiency (>90%) in just 4 days, reducing the need for additional purification or enrichment steps. Hence, our method could not only be cost-effective but also resource-saving when manufacturing ECs for regenerative medicine. In addition, it is important to note that modified mRNA vectors are non-viral, non-integrating, and inherently transient, which from a translational standpoint is also beneficial. We anticipate our protocol could have broad application in regenerative medicine because it provides a reliable means to obtain autologous h-iECs for vascular therapies. The advent of high-throughput electroporation systems, as well as advances in scalability and clinical grade materials, should enable further refinements to this protocol for the generation of clinical-grade h-iECs.

## MATERIALS AND METHODS

### Isolation and culture of human MSCs, ECFCs and uEPs

Human MSCs (h-MSCs) were isolated from the white adipose tissue as previously described (*34*). h-MSCs were cultured on uncoated plates using MSC-medium: MSCGM (Lonza, Cat No. PT-3001) supplemented with 10% GenClone FBS (Genesee, Cat No. 25-514), 1 x penicillin-streptomycin-glutamine (PSG, ThermoFisher, Cat No. 10378106). All experiments were carried out with h-MSCs between passage 6-10. Human ECFCs were isolated from umbilical cord blood samples in accordance with an Institutional Review Board-approved protocol as previously described (*35*). All ECFCs were derived from umbilical cord blood samples obtained from normal term deliveries. Samples were collected *ex utero* using heparinized tubes. ECFCs were cultured on 1% gelatin-coated plates using ECFC-medium: EGM-2 (except for hydrocortisone; PromoCell, Cat No. C22111) supplemented with 10% FBS, 1x PSG. All experiments were carried out with ECFCs between passage 6-8. Human urine-derived epithelial cells (uEPs) were isolated from urine samples and were cultured on 1% gelatin-coated plates using ECFC-medium. All experiments were carried out with uEPs up to passage 4.

### Generation and culture of human iPSCs

Human induced pluripotent stem cells (h-iPSCs) were generated via non-integrating episomal transferring of selected reprogramming factors (Oct4, Sox2, Klf4, L-Myc, Lin28). Briefly, four plasmids encoding h-oct4, h-sox2, h-klf4, h-myc, h-lin-28 and EBNA-1 (Addgene plasmids #27077, #27078, #27080, and #37624 deposited by Shinya Yamanaka) were introduced via electroporation into h-MSCs, ECFCs and uEPs. Transfected cells were then cultured with mTeSR-E7 medium (STEMCELL, Cat No. 05910). H-iPSC colonies spontaneously emerged between days 15-25. Colonies were then picked and transferred to a Matrigel-coated (Corning, Cat No. 354277), feeder-free culture plate for expansion and were routinely checked for absence of mycoplasma. H-iPSCs were cultured in mTeSR1 medium (STEMCELL, Cat No. 85850) on 6-well plates coated with Matrigel. At 80% confluency, h-iPSCs were detached using TrypLE select (ThermoFisher, Cat No. 12563-029) and split at a 1:6 ratio. Culture media were changed daily. h-iPSCs phenotype was validated by expression of pluripotent transcription factors OCT4, NANOG, and SOX2 and by the ability to form teratomas. Teratoma formation assay was performed by injecting 1 million h-iPSCs mixed in 100 μL Matrigel into the dorsal flank of nude mice (Jackson Lab). Four weeks after the injection, tumors were surgically dissected from the mice, weighed, fixed in 4% formaldehyde, and embedded in paraffin for histology. Sections were stained with hematoxylin and eosin (H&E).

### Electroporation

Electroporation was routinely used to introduce plasmids, modified mRNA and proteins into the cells as described for each experiment. Electroporation was carried out with a Neon electroporation system (ThermoFisher). Unless specified otherwise, electroporation parameters were set as 1150 v for pulse voltage, 30 ms for pulse width, 2 for pulse number, 3 mL of electrolytic buffer and 100 μL resuspension buffer R in 100 μL reaction tips (ThermoFisher, Cat No. MPK10096).

### Establishment of *KDR* and *ETV2* knock out h-iPSC lines

Alt-R™ CRISPR-Cas9 system (Integrated DNA Technologies, IDT) was used to knock out *KDR* (gRNA: ACGGACTGTACCATTTCGTG) and *ETV2* (gRNA: GAGCCTACAAGTGCTTCTAC) in h-iPSCs. Briefly, guide RNA (gRNA) was prepared by mixing crRNA and tracrRNA (IDT, Cat No.1072533) to a final duplex concentration of 40 μΜ. Ribonucleoprotein (RNP) complex was prepared with 1 μL volume of 61 μΜ Cas9 protein (IDT, Cat No. 1074181) complexed with 2.5 μL of gRNA for 15 min at room temperature. Following incubation, RNP complexes were diluted with 100 μL R buffer and mixed with one million pelleted h-iPSCs for electroporation. Two days later, h-iPSCs were dissociated into single cells and plated at 2,000 cells per 10 cm dish in mTeSR1 supplemented with CloneR (STEMCELL, Cat No. 5888). Single cells were able to grow and form single visible colonies after 10 days. 48 colonies were randomly picked based on morphology and were then mechanically disaggregated and replated into individual wells of 48-well plates. Colonies were then expanded in culture as described above. To validate the knock out genes in each clone, genomic DNA templates were prepared by lysing cells in QuickExtract DNA extraction solution (Lucigen, Cat No. QE0905T). Target regions were amplified by using specific PCR primers (ETV2_F: CACTCGGGATCCGTTACTCC; ETV2_R: GTTCGGAGCAAACGGTGAGA, KDR_F: CAAGCCCTTTGTTGTACTCAATTCT; KDR_R: ATTAATTTTTCAGGGGACAGAGGGA) and KAPA HiFi HotStart PCR kit (KAPA Biosystems, Cat No. KK2601). Sanger sequencing (Genewiz) was performed to identify mutant clones.

### Establishment of h-iPSC line expressing GFP

h-iPSCs were dissociated and filtered through 40 μm cell strainer to get single cells. For electroporation, 1 million h-iPSCs were resuspended in 100 μL buffer mixed with 2 μg PB-EF1A-GFP-puro plasmid (VectorBuilder) and 1 μg transposase plasmid (VectorBuilder). The electroporated cells were then plated on a 35-mm Matrigel-coated dish in mTeSR1 medium with 10 μM Y27632 (Selleckchem, Cat No. S1049). After 48 hours, culture medium was replaced by mTeSR1 medium with 10 μg/mL puromycin (Sigma, Cat No. P8833) and changed daily for 3 to 4 days.

### Modified mRNA synthesis and formulation

Chemically modified mRNA encoding ETV2 (modRNA(ETV2)) was generated by TriLink BioTechnologies, LLC. In brief, modRNA(ETV2) was synthesized *in vitro* by T7 RNA polymerase-mediated transcription from a linearized DNA template, which incorporates the 5’ and 3’ UTRs and a poly-A tail. Specifically, ETV2 cDNA was cloned into the mRNA expression vector pmRNA, which contains a T7 RNA polymerase promoter, an unstructured synthetic 5’ UTR, a multiple cloning site, and a 3’ UTR that was derived from the mouse *α*-globin 3’ gene. *In vitro* transcriptional (IVT) reaction (1 mL-scale) was performed to generate unmodified mRNA transcripts with wild type bases and a poly-A tail. Co-transcriptional capping with CleanCap Cap1 AG trimer yields a naturally occurring Cap1 structure. DNase treatment was used to remove DNA template. 5’-triphosphate were removed by phosphatase treatment to reduce innate immune response. After elution through silica membrane, the purified RNA was dissolved in RNase-free sodium citrate buffer (1 mM, pH 6.4).

### Differentiation of h-iPSCs into h-iECs

<u>S1-modETV2 protocol</u> (4 days) – h-iPSCs were dissociated into single cells with TrypLE select (ThermoFisher, Cat No. 12563-029) and plated on Matrigel at a density of 50,000 cells/cm^2^ in mTeSR1 medium with 10 μM Y27632. After 24 h, the medium was changed to S1 medium consisting of basal medium supplemented with 6 μM CHIR99021 (Sigma, Cat No. SML1046). Basal medium was prepared by adding 1x GlutaMax supplement (ThermoFisher, Cat No. 35050061) and 60 μg/mL L-Ascorbic acid (Sigma, Cat No. A8960) into Advanced DMEM/F12 (ThermoFisher, Cat No. 12634010). After 48 h, h-MPCs were dissociated into single cells and then transfected with modRNA(ETV2) by either electroporation or lipofection. For electroporation, 2 million h-MPCs were resuspended in 100 μL buffer mixed with 1 μg modETV2. Electroporated cells were then seeded on a 60-mm Matrigel-coated dish in modETV2 medium consisting of basal medium supplemented with 50 ng/mL VEGF-A (PeproTech, Cat No. 100-20), 50 ng/mL FGF-2 (PeproTech, Cat No. 100-18B), 10 ng/mL EGF (PeproTech, Cat No. AF-100-15) and 10 μM SB431542 (Selleckchem, Cat No. S1067). For lipofection, 3 μL lipofectamine RNAiMax (ThermoFisher, Cat No. 13778030) were diluted in 50 μL Opti-MEM (ThermoFisher, Cat No. 31985062) and 0.6 μg modRNA(ETV2) diluted in another 50 μL Opti-MEM. Lipofectamine and modRNA(ETV2) were then mixed and incubated for 15 min at room temperature. The lipid/RNA complex was added to 0.5 million h-MPCs in modETV2 medium and transfected cells were then seeded on a 35-mm Matrigel-coated dish. Upon transfection (electroporation or lipofection), cells were cultured for another 48 h before purification. Medium was changed every day throughout this protocol. ModRNA encoding GFP (TriLink, Cat No. L-7601) at a concentration of 0.2 μg per million h-MPCs served as negative control.

<u>Early modETV2 protocol</u> (2 days) – h-iPSCs were dissociated into single cells and then transfected with modRNA(ETV2) by electroporation. For electroporation, 2 million h-iPSCs were resuspended in 100 μL buffer and mixed with 1.5 μg modRNA(ETV2). Electroporated cells were then plated on a 60-mm Matrigel-coated dish in mTeSR1 medium with 10 μM Y27632. After 24 h, the medium was changed to mTeSR1 medium with 10 μM SB431542 for another 24 h.

<u>S1-S2, method #1</u> (4 days) – h-iPSCs were dissociated into single cells with TrypLE select and plated on Matrigel at a density of 50,000 cells/cm^2^ in mTeSR1 medium with 10 μM Y27632. After 24 h, the medium was changed to S1 medium consisting of basal medium supplemented with 6 μM CHIR99021. Basal medium was prepared by adding 1x GlutaMax supplement and 60 μg/mL L-Ascorbic acid into Advanced DMEM/F12. After 48 h, the differentiation medium was changed to S2 medium for 48 h. S2 medium consisted of basal medium supplemented with 50 ng/mL VEGF-A, 50 ng/mL FGF-2, 10 ng/mL EGF and 10 μM SB431542. Medium was changed every day throughout this protocol.

<u>S1-S2, method #2</u> (8 days) – h-iPSCs were dissociated into single cells with TrypLE select and plated on Matrigel at a density of 60,000 cells/cm^2^ in mTeSR1 medium with 10 μM Y27632. After 24 h, the medium was changed to STEMdiff APEL2 medium (STEMCELL, Cat No. 05275) supplemented with 6 μM CHIR99021. After 48 h, the differentiation medium was changed to S2 medium for 48 h. S2 medium consisted of STEMdiff APEL2 medium supplemented with 50 ng/mL VEGF-A, 10 ng/mL FGF-2, and 25 ng/mL BMP4 (PeproTech, Cat No. 120-05ET). Medium was changed every day throughout the first 4 days. Cells were lifted at day 4 and seeded on p100 dish at 50,000 cells/cm^2^ in EGM2 with additional 50 ng/mL VEGF for another 4 days. This protocol is adapted from Harding *et al*. 2017 (*18*).

<u>S1-S2, method #3</u> (5 days) – h-iPSCs were dissociated into single cells with TrypLE select and plated on Matrigel at a density of 50,000 cells/cm^2^ in mTeSR1 medium with 10 μM Y27632. After 24 h, the medium was changed to basal medium supplemented with 1 μM CP21R7 (Selleckchem, Cat No. S7954) and 20 ng/mL BMP4. Basal medium was prepared by adding 1x B27 supplement (ThermoFisher, Cat No. 17504044) and 1x N2 (ThermoFisher, Cat No. 17502048) into DMEM/F12 (ThermoFisher, Cat No. 11330032). After 72 h, the differentiation medium was changed to S2 medium for 48 h. S2 medium consisted of StemPro-34 SFM (ThermoFisher, Cat No. 10639011) supplemented with 50 ng/mL VEGF-A, and 10 μM DAPT (Selleckchem, Cat No. S2215). Medium was changed every day throughout this protocol. This protocol is adapted from Sahara *et al*. 2014 (*19*).

<u>S1-S2, method #4</u> (5 days) – h-iPSCs were dissociated into single cells with TrypLE select and plated on Matrigel at a density of 50,000 cells/cm^2^ in mTeSR1 medium with 10 μM Y27632. After 24 h, the medium was changed to basal medium supplemented with 8 μM CHIR99021. Basal medium was prepared by adding 1x B27 supplement and 1x N2 into DMEM/F12. After 72 h, the differentiation medium was changed to S2 medium for 48 h. S2 medium consisted of StemPro-34 SFM supplemented with 200 ng/mL VEGF-A, and 2 μM forskolin (Sigma, Cat No. F3917). Medium was changed every day throughout this protocol. This protocol is adapted from Patsch *et al*. 2015 (*4*).

### Purification and expansion of h-iECs

At indicated time points after differentiation, h-iECs were dissociated into single cells and sorted into CD31+ and CD31-cells using magnetic beads coated with anti-human CD31 antibodies (DynaBeads, ThermoFisher, Cat No. 11155D). The purified CD31+ h-iECs were then expanded in culture on 10-cm dishes coated with 1% gelatin. Culture medium for h-iECs was prepared by adding Endothelial Cell Growth medium 2 kit supplements into basal medium (except for hydrocortisone, PromoCell, Cat No. C22111) with 1x GlutaMax supplement and 10 μM SB431542.

### RNA-Seq analysis

The following groups were analysed: h-iPSCs, human ECFCs, and h-iECs generated with three protocols: S1-S2, S1-modETV2, and early modETV2. Each group consists of 3 biological replicates. Total RNA from h-iECs which have been expanded for 7 days was extracted using Rneasy Mini Kit (Qiagen) following the manufacturer’s protocol. RNA quantity and quality were checked with nanodrop and Agilent Bioanalyzer instrument. Libraries were prepared and sequenced by GENEWIZ (NJ, USA). Library preparation involved mRNA enrichment and fragmentation, chemical fragmentation, first and second strand cDNA synthesis, end repair and 5’ phosphorylation, Da-tailing, adaptor ligation and PCR enrichment. The libraries were then sequenced using Illumina HiSeq2500 platform (Illumina, CA) using 2×150 paired end configuration. The raw sequencing data (FASTQ files) was examined for library generation and sequencing quality using FastQC (https://www.bioinformatics.babraham.ac.uk/projects/fastqc/) to ensure data quality was suitable for further analysis. Reads were aligned to UCSC hg38 genome using the STAR aligner (*36*). Alignments were checked for evenness of coverage, rRNA content, genomic context of alignments, complexity, and other quality checks using a combination of FastQC, Qualimap (*37*) and MultiQC (*38*). The expression of the transcripts was quantified against the Ensembl release GRCh38 transcriptome annotation using Salmon. These transcript abundances were then imported into R (version 3.5.1) and aggregated to the gene level with tximport. Differential expression at the gene level was called with DESeq2 (*39*). Pairwise differential expression analysis between groups was performed using Wald significance test. The P values was corrected for multiple hypothesis testing with the Benjamini-Hochberg false-discovery rate procedure (adjusted P value). Genes with adjusted P value < 0.05 were considered significantly different. Hierarchical clustering and PCA analysis were performed on DESeq2 normalized, rlog variance stabilized reads. All samples comparison was performed using Likelihood Ratio Test (LRT). Heat maps of the differential expressed genes and enriched gene sets were generated with pheatmap package. Functional enrichment of differential expressed genes, using gene sets from Gene Ontology (GO), was determined with Fisher’s exact test as implemented in the clusterProfiler package. The RNA-Seq datasets are deposited online with SRA accession number: PRJNA509218.

### Flow cytometry

Cells were dissociated into single-cell suspensions using TrypLE and washed with PBS supplemented with 1% BSA and 0.2 mM EDTA. In indicated experiments, cells were stained with flow cytometry antibodies and analyzed using a Guava easyCyte 6HT/2L flow cytometer (Millipore Corporation, Billerica, MA) and FlowJo software (Tree Star Inc., Ashland, OR). Antibody labeling was carried out for 10 min on ice followed by 3 washes with PBS buffer. Antibody information is detailed in Supplementary Table 1.

### Microscopy

Images were taken using an Axio Observer Z1 inverted microscope (Carl Zeiss) and AxioVision Rel. 4.8 software. Fluorescent images were taken with an ApoTome.2 Optical sectioning system (Carl Zeiss) and 20x objective lens. Non-fluorescent images were taken with an AxioCam MRc5 camera using a 5x or 10x objective lens.

### Immunofluorescence staining

Cells were seeded in 8-well LAB-TEK chamber slides at a density of 60,000 cells/cm^2^. After confluency, cells were fixed in 4% paraformaldehyde (PFA), permeabilized with 0.1% Triton X-100 in PBS, and then blocked for 30 min in 5% horse serum (Vector, Cat No. S-2000). Subsequently, cells were incubated with primary antibodies for 30 min at room temperature (RT). Cells were washed 3 times with PBS and then incubated with secondary antibodies for 30 min at RT. Cells were washed 3 times with PBS and stained with 0.5 μg/mL DAPI for 5 min. Slides were mounted with DAKO fluorescence mounting medium (Agilent, Cat No. S302380-2). Antibody information is detailed in Supplementary Table 1.

### Spheroid sprouting assay

EC spheroids were generated by carefully depositing 500 h-MSCs and 500 h-iECs-GFP in 20 μL spheroid-forming medium on the inner side of a 10-cm dish lid. The spheroid-forming medium contained 0.24% (w/v) methyl cellulose (Sigma, Cat No. M0512). The lid was then turned upside down and placed on top of the plate filled with 10 mL sterile water. EC spheroids were collected after 2 days in culture and embedded in fibrin gel prepared with 5 mg/mL fibrinogen (Sigma, Cat No. F8630) and 0.5 U/mL thrombin (Sigma, Cat No. T-9549). A 100 μL-fibrin gel/spheroid solution was spotted into the center of a 35-mm glass bottom dish (MatTek, Cat No. P35G-1.5-10-C) and incubated for 10 mins at 37 °C for solidification. Gel/spheroid constructs were kept in culture for 3 days. GFP+ sprouts were imaged using an inverted fluorescence microscope and sprout lengths were measured by ImageJ.

### Shear stress response assay

Confluent monolayers of h-iECs in a 100-mm culture dish were subjected to orbital shear stress for 24 h at a rotating frequency of 150 rpm using an orbital shaker (VWR, Model 1000) positioned inside a cell culture incubator. After 24 h, cells were fixed in 4% PFA and stained using an anti-human VE-Cadherin antibody. Alignment of ECs was visualized using an inverted fluorescence microscope under a 10X objective. Only the cells in the periphery of the culture dish were imaged. Cell orientation angles were measured by ImageJ.

### Nitric oxide (NO) production assay

Cells were cultured on gelatin-coated 12-well plates (2×10^5^ cells per well) in h-iECs media. To measure nitric oxide (NO), media were changed to fresh media containing 1 μM DAF-FM (Cayman, Cat No. 18767). Cells were cultured for 30 min and then harvested for flow cytometric analysis and fluorescent imaging. In order to suppress NO production, h-iECs were cultured in the presence of 5 mM L-NAME (Cayman, Cat No. 80210) for 24 h. DAF-FM is nonfluorescent until it reacts with NO to form a fluorescent benzotriazole (FITC channel). The mean fluorescence intensities (MFIs) were measured by calculating the geometric mean in FlowJo.

### Leukocyte adhesion molecules and leukocyte adhesion assays

Cells were cultured on a gelatin-coated 48-well plate (10^5^ cells per well) in h-iEC medium. At confluency, cells were treated with or without 10 ng/mL TNF-α (Peprotech, Cat No. 300-01A) for 5 h. Cells were then lifted and treated with anti-ICAM1, anti-E-selectin or anti-VCAM1 antibodies for flow cytometry. For leukocyte adhesion assay, human HL-60 leukocytes were used. HL-60 cells were culture in leukocyte medium consisting of RPMI-1640 (ThermoFisher, Cat No. 11875093) supplemented with 20% FBS. 2 x 10^5^ HL-60 cells were suspended in 0.2 mL fresh leukocyte medium and added to each well. After gentle shaking for 45 min in cold room, plates were gently washed twice with cold leukocyte media. Cells were fixed in 2.5%(v/v) glutaraldehyde at RT for 30 min and then imaged. Bound leukocytes were quantified by ImageJ analysis software.

### Smooth muscle cell differentiation assay

2×10^4^ h-MSCs and 5×10^4^ h-iECs were plated in one well of 8-well LAB-TEK chamber slide coated by 1% gelatin and cultured in h-iECs medium without SB431542 for 7 days. Smooth muscle cell positive cells were stained with an anti-smooth muscle myosin heavy chain 11 antibody and ECs and nucleus were stained by anti-VECAD antibody and DAPI, respectively. h-MSCs that were transduced with lentivirus to express GFP (h-MSCs-GFP) were used in indicated experiments. Antibody information is detailed in Supplementary Table 1.

### Tube formation assay

8×10^3^ h-iECs were plated in one well of 96-well plate on top of solidified Matrigel (50 μL) with h-iECs media. After 6 h, cells were incubated with 1 μM Calcein-AM (Biolegend, Cat No. 425201) for 10 min and then imaged using a fluorescence microscope. Numbers of branches were counted by ImageJ.

### *In vivo* vascular network-forming assay

Six-week-old NOD/SCID mice were purchased from Jackson Lab (Boston, MA). Mice were housed in compliance with Boston Children’s Hospital guidelines, and all animal-related protocols were approved by the Institutional Animal Care and Use Committee. Human iECs for implantation were expanded for 7 days *in vitro* after differentiation unless otherwise specified. H-iECs were pretreated with 20 μM caspase inhibitor/Z-VAD-FMK (APExBio, Cat No. A1902) and 0.5 μM BCL-XL-BH4 (Millipore, Cat No. 197217) in h-iECs medium overnight before implantation. Briefly, h-iECs and h-MSCs (2×10^6^ total per mice, 1:1 ratio) or h-iECs alone (1×10^6^ cells per mice) were resuspended in 200 μL of pH neutral pre-gel solution containing 3 mg/mL of bovine collagen I (Trevigen, Cat No. 3442-050-01), 3 mg/mL of fibrinogen, 50 μL Matrigel (Corning, Cat No. 354234), 1 μg/mL of FGF2 (Peprotech, Cat No. 100-18B) and 1 μg/mL EPO (ProSpec, Cat No. CYT-201). During anesthesia, mice were firstly injected with 50 μL of 10 U/mL thrombin (Sigma, Cat No. T4648) subcutaneously and then injected with 200 μL cell-laden pre-gel solution into the same site. All experiments were carried out in 5 mice and explants were harvested after 1 week and 1 month.

### Histology and immunofluorescence staining

Explanted grafts were fixed overnight in 10% buffered formalin, embedded in paraffin and sectioned (7-µm-thick). Microvessel density was reported as the average number of erythrocyte-filled vessels (vessels/mm^2^) in H&E-stained sections from the middle of the implants as previously were deparaffinized and antigen retrieval was carried out with boiling citric buffer (10 mM sodium citrate, 0.05% Tween 20, pH 6.0) for 30 min. Proteinase K (Abcam, Cat No. ab64220) treatment for 15 min was only applied to retrieve the mouse-specific CD31 described. For immunostaining, sections antigen. Sections were then blocked for 30 min in 5% horse serum and incubated with primary and secondary antibodies for 30 min at RT. Fluorescent staining was performed using fluorescently-conjugated secondary antibodies followed by DAPI counterstaining. Human-specific anti-CD31 antibody and UEA-1 lectin were used to stain human blood vessels. Perivascular cells were stained by anti-alpha smooth muscle actin antibody. Primary and secondary antibodies are detailed in Supplementary Table 1. The Click-It Plus TUNEL assay (ThermoFisher, Cat No. C10617) was used to detect apoptotic cells in tissue.

### Perfusion of human vessels

To test for perfusion of the human vessels, one week after h-iECs were implanted, biotinylated UEA-I (50 μg per mouse) was injected through the retro-orbital sinus. 30 min after injection, mice were euthanized, and the implants were harvested. Perfused human vessels were identified as lumens with bound biotinylated UEA-I. Streptavidin Texas Red was applied to the histological sections of the grafts to visualize UEA-1+ vessels. Sections were also stained with human-specific anti-CD31 to confirm the presence of human endothelial cells.

### Western Blot

Cells were lysed with RIPA Lysis Buffer (Thermo Fisher, Cat No. 89901) in the presence of protease inhibitor. The concentration of extracted protein was measured using Pierce 660 nm Protein Assay Reagent (Thermo Fisher, Cat No. 1861426) and Bio-Rad SmartSpecTM 3000 spectrophotometer. 40 μg of whole cell protein lysate were applied to 4%-15% Mini-PROTEAN TGX precast protein Gel (Bio-Rad, Cat No. 4561084) with electrophoresis and then transferred to a nitrocellulose membrane. The probed primary antibodies were detected by using Horseradish Peroxidase (HRP)-conjugated secondary antibodies and the Enhanced Chemiluminescent (ECL) detection system (GE Health, Cat No. 28906836). Primary and secondary antibodies are detailed in Supplementary Table 1.

### Quantitative RT-PCR

Quantitative RT-PCR (qRT-PCR) was carried out in RNA lysates prepared from cells in culture. Total RNA was isolated with a RNeasy kit (Qiagen, Cat No. 74106) and cDNA was prepared using reverse transcriptase III (ThermoFisher, Cat No. 4368814), according to the manufacturer’s instructions. Quantitative PCR was performed using SRBR Green Master Mix (ThermoFisher, Cat No. A25776), and detection was achieved using the StepOnePlus Real-time PCR system thermocycler (Applied Biosystems). Expression of target genes was normalized to GAPDH. Real-time PCR primer sequences are listed in Supplementary Table 2.

### Statistical analyses

Unless otherwise stated, data were expressed as mean ± standard deviation of the mean (s.d.). For comparisons between two groups, means were compared using unpaired two-tailed Student’s t-tests. Comparisons between multiple groups were performed by ANOVA followed by Bonferroni’s post-test analysis. Samples size, including number of mice per group, was chosen to ensure adequate power and were based on historical data. No exclusion criteria were applied for all analyses. All statistical analyses were performed using GraphPad Prism v.5 software (GraphPad Software Inc.). P<0.05 was considered statistically significant.

## Acknowledgments

Histology was supported by Core Facility of the Dana-Farber/Harvard Cancer Center (P30 CA06516).

## Author contributions

K.W., R.-Z.L. and J.M.M.-M. conceived and designed the project. K.W., R.-Z.L., A.H.N., X.H., C.-N.L., J.N., G.W., X.W., and J.M.M.-M. performed the experimental work. All authors discussed and analyzed the data and edited the results. A.H.N., G.W., W.T.P., and G.M.C. provided crucial material. K.W. and J.M.M.-M. wrote the manuscript.

## Data and materials availability

All data needed to evaluate the conclusions in the paper are present in the paper and/or the Supplementary Materials. Additional data related to this paper may be requested from the authors.

## SUPPLEMENTARY MATERIAL

**Fig. S1.**
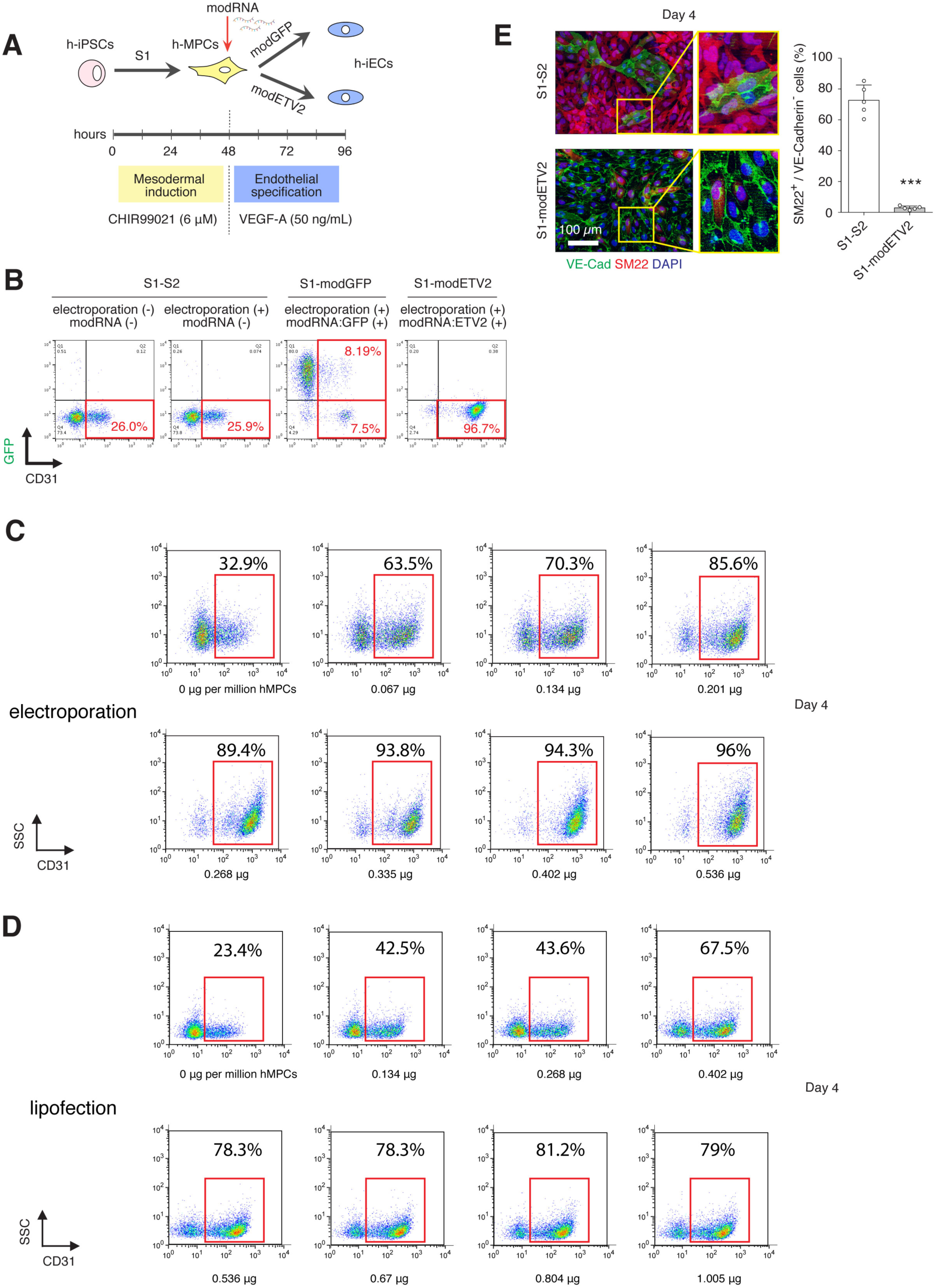
Derivation of h-iECs from h-iPSCs with ETV2 modRNA. (**A**) Schematic of optimized two-stage endothelial differentiation protocol. Stage 1: conversion of h-iPSCs into h-MPCs mediated by the GSK-3 inhibitor CHIR99021. Stage 2: transfection of h-MPCs with modRNA encoding *ETV2* and culture in chemically defined medium. A group that used modRNA:GFP served as control. (**B**) Conversion efficiency of h-iPSCs into VE-Cadherin+/CD31+ h-iECs measured by flow cytometry at day 4 for both the S1-S2 (left; no electroporation, no modRNA) and S1-modETV2 (right) protocols. Groups corresponding to electroporation without modRNA and electroporation with modRNA encoding GFP served as controls for the S1-modETV2 group. (**C-D**) Effect of modRNA(ETV2) concentration on h-iPSC-to-h-iEC conversion efficiency at 96 h using the S1-modETV2 differentiation protocol. (**C**) Titration analysis by flow cytometry for electroporation-based delivery of modRNA. (**D**) Titration analysis by flow cytometry for lipofection-based delivery of modRNA. **(E)** Immunofluorescence staining for VE-Cadherin and SM22 between S1-S2 and S1-modETV2 protocols at day 4. Nuclei stained by DAPI. Scale bar, 100 μm. Percentage of SM22+/VE-Cadherin-cells. Bars represent mean ± s.d.; n = 5. \*\*\**P* < 0.001 between S1-S2 and S1-modETV2 protocols.

**Fig. S2.**
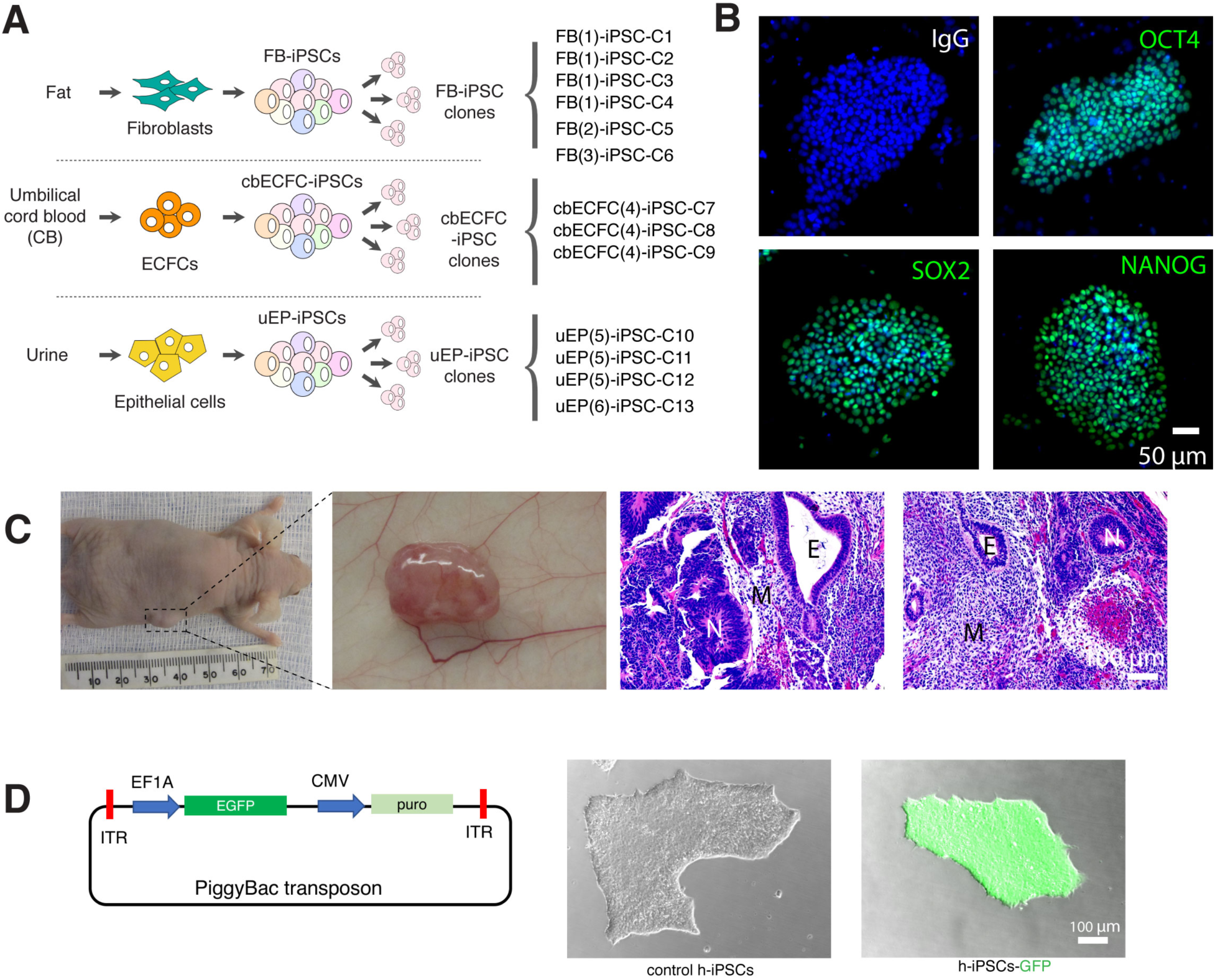
Characterization of h-iPSC clones generated from different donors and tissues. (**A**) Schematic of generation of thirteen h-iPSC clones from dermal fibroblasts (dFB), umbilical cord blood-derived ECFCs (cbECFC), and urine-derived epithelial cells (uEP). (**B**) Immunofluorescence staining for pluripotency markers including OCT4, SOX2 and NANOG. Nuclei stained by DAPI. Scale bar, 50 μm. (**C**) Teratoma formation upon implantation of h-iPSCs into nude mice for 4 weeks. Hematoxylin and eosin (H&E) staining of explanted tumors showed three germ layers including neuroepithelial rosettes (N), endodermal gut-like tissues (E) and mesenchymal stromal tissue (M). Scale bar, 100 μm. (**D**) Generation of EGFP-labeled h-iPSC with transposase mediated knock-in of a PiggyBac transposon plasmid. After puromycin selection and clonal expansion, homogenous green fluorescent clones of h-iPSC-GFP were generated. Scale bar, 100 μm.

**Fig. S3.**
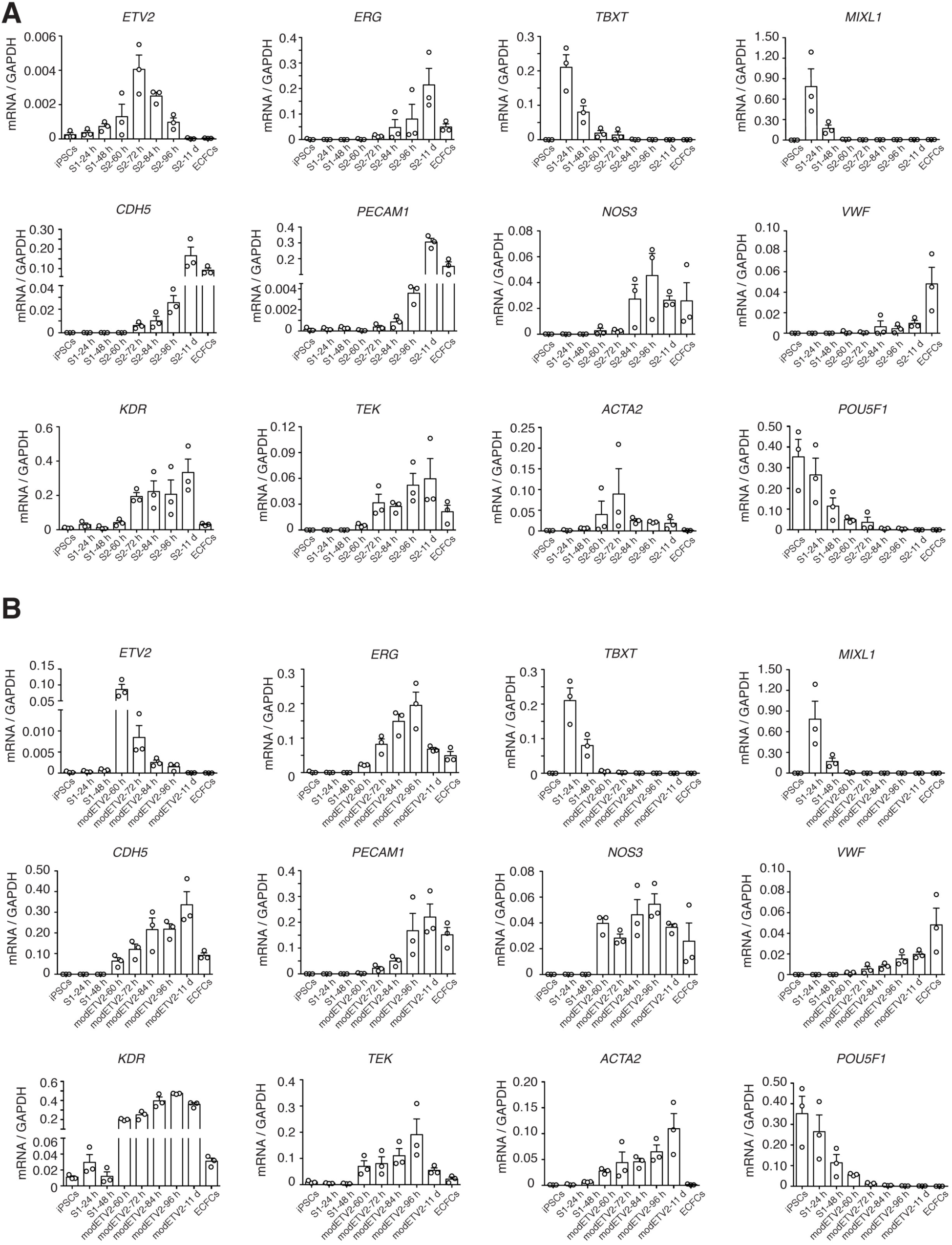
Time course analysis of mRNA expression (qRT–PCR). Genes including mesodermal markers (*TBXT* and *MIXL1*), endothelial commitment transcription factors (*ETV2* and *ERG*), endothelial markers (*PECAM1, CDH5, NOS3, VWF, TEK* and *KDR*), pluripotency marker (*POU5F1*) and smooth muscle marker (*ACTA2*) in the standard S1-S2 protocol (**A**) the S1-modETV2 protocol (**B**). Data normalized to *GAPDH* expression.

**Fig. S4.**
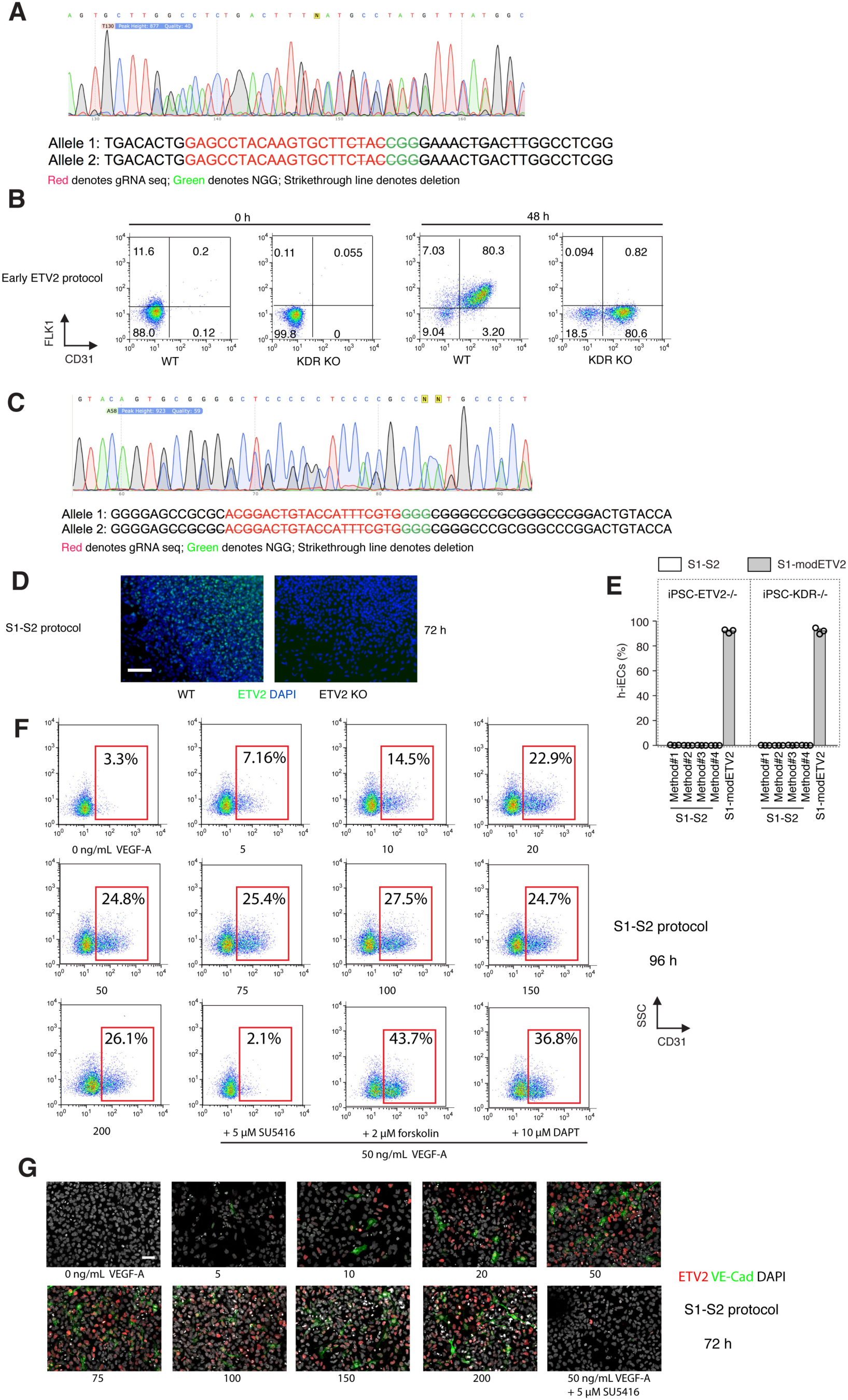
VEGF signaling and activation of endogenous ETV2 in the standard S1-S2 differentiation protocol. (**A-D**) Generation of h-iPSCs-KDR^-/-^ and h-iPSCs-ETV2^-/-^ clones by CRISPR/Cas9. (**A**) Sanger sequencing of the two edited alleles encoding the 3^rd^ exon of *KDR*. (**B**) Flow cytometry showed the conversion of h-iPSCs-KDR^-/-^ into FLK1-/CD31+ h-iECs at 48 h using the early modETV2 protocol. (**C**) Sanger sequencing of the two edited alleles encoding the 4^th^ exon of *ETV2*. (**D**) Immunofluorescence staining for ETV2 at 72 h using the S1-S2 differentiation protocol. Nuclei stained by DAPI. Scale bar, 200 μm. (**E**) Differences in differentiation efficiency between four alternative S1-S2 methodologies and the S1-modETV2 protocol for h-iPSC clones lacking either ETV2 and KDR (h-iPSC-ETV2^-/-^ and h-iPSC-KDR^-/-^). Only S1-modETV2 protocol could successfully derive h-iECs from either h-iPSC-ETV2^-/-^ or h-iPSC-KDR^-/-^ cell lines with high efficiency. In contrast, the four alternative S1-S2 methodologies failed to get any h-iECs. (**F-G**) Effect of VEGF-A concentration on h-iEC yield using the S1-S2 differentiation protocol. (**F**) Dose dependent conversion efficiency of h-iPSCs into CD31+ h-iECs by flow cytometry. (**G**) Immunofluorescence staining for ETV2 and VE-Cadherin at 72 h with different concentrations of VEGF-A. Nuclei stained by DAPI. Scale bar, 50 μm.

**Fig. S5.**
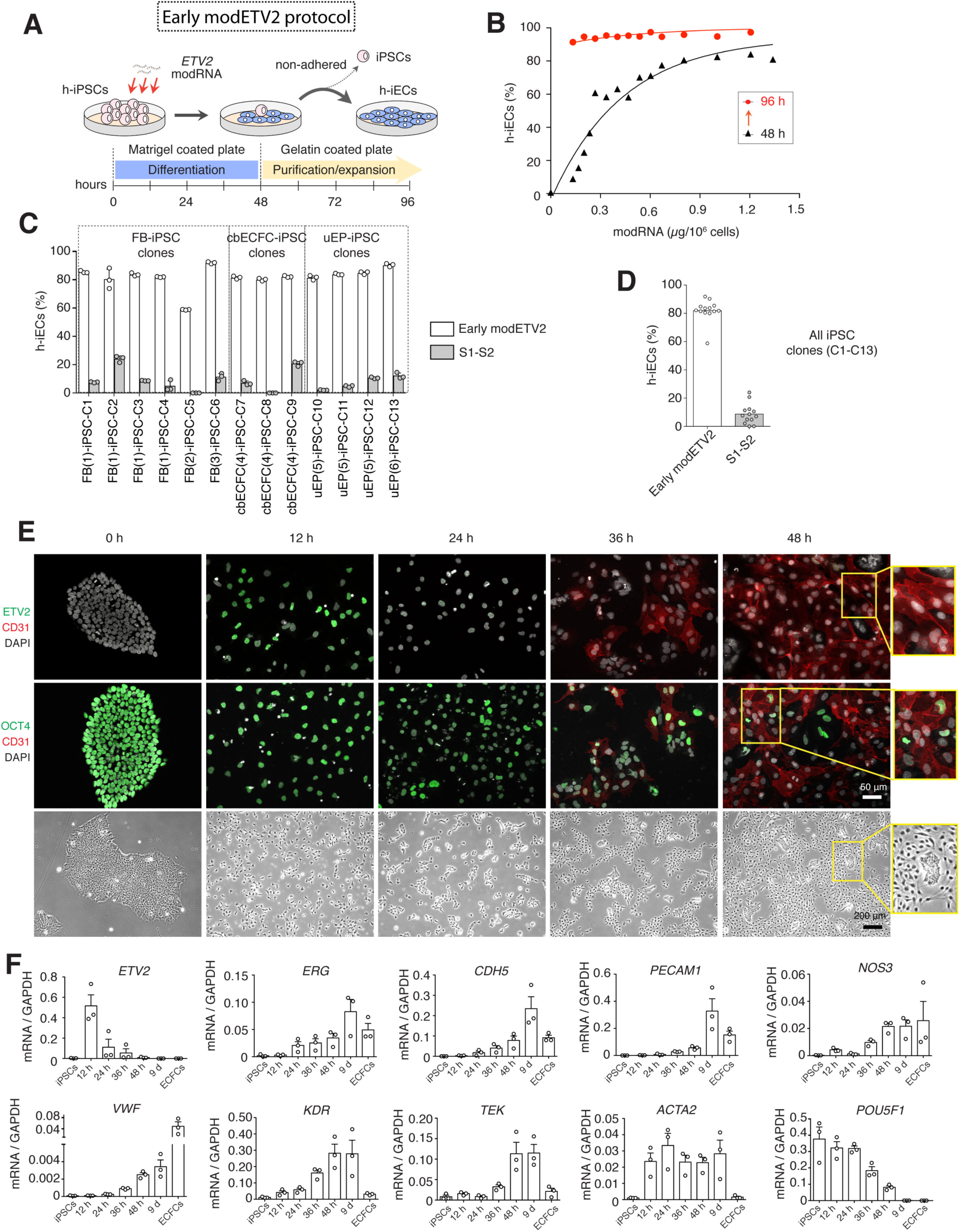
Generation of h-iECs with the early modETV2 differentiation protocol. (**A**) Schematic of the early modETV2 protocol. (**B**) Effect of modRNA(ETV2) concentration on h-iPSC-to-h-iEC conversion efficiency at 48 h and 96 h. Titration analysis by flow cytometry for electroporation-based delivery of modRNA. (**C**) Flow cytometry analysis of differentiation efficiency at 48 h (early modETV2 protocol) and 96 h (S1-S2 protocol) in 13 h-iPSC clones generated from dermal fibroblasts (dFB), umbilical cord blood-derived ECFCs (cbECFC), and urine-derived epithelial cells (uEP). (**D**) Differences in differentiation efficiency between early modETV2 and S1-S2 protocols for all 13 h-iPSC clones. Bars represent mean ± s.d. (**E**) Time course immunofluorescence staining for ETV2, CD31 and OCT4 during the early modETV2 differentiation protocol. Nuclei stained by DAPI. Scale bar, 100 μm. Phase contrast micrographs represent time course morphological changes of cells during the early modETV2 protocol. Scale bar, 200 μm. (**F**) Time course analysis of mRNA expression (qRT–PCR) of mesodermal markers (*TBXT* and *MIXL1*), endothelial commitment transcription factors (*ETV2* and *ERG*), endothelial markers (*PECAM1, CDH5, NOS3, VWF, TEK* and *KDR*), pluripotency marker (*POU5F1*) and smooth muscle marker (*ACTA2*) in h-iECs generated with the early modETV2 protocol. Data normalized to *GAPDH* expression.

**Fig. S6.**
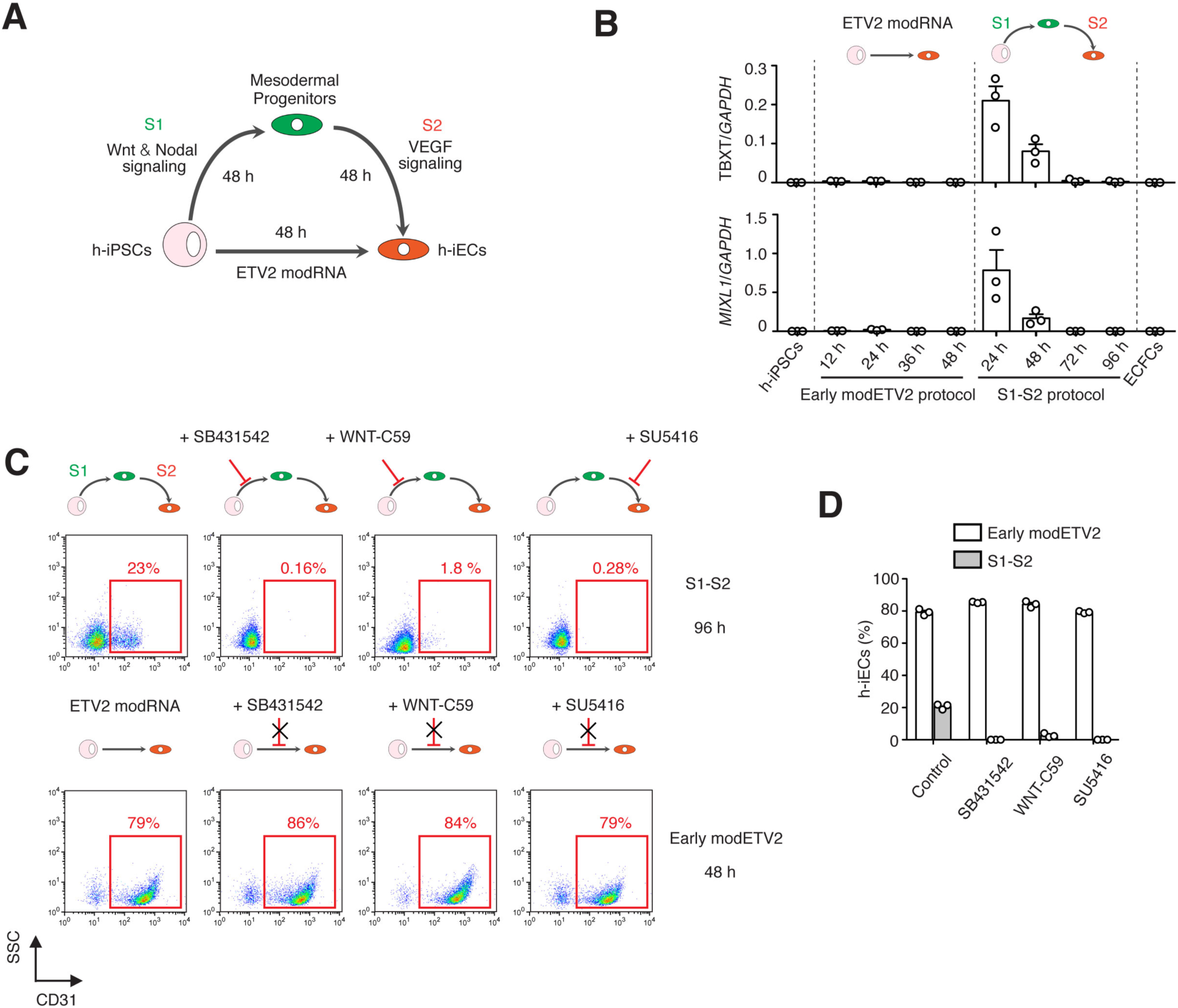
Early modETV2 protocol bypasses the intermediate mesodermal stage. (**A**) Schematic of S1-S2 and early modETV2 differentiation protocols. (**B**) Time course analysis of mRNA expression (qRT–PCR) of mesodermal markers (*TBXT* and *MIXL1*) in the S1-S2 and the early modETV2 protocols. (**C**) Effect of small molecule inhibitors SB431542 (10 μM), Wnt-C59 (5 μM), and SU5416 (5 μM) on the percentage of h-iECs generated with the S1-S2 (96 h) and the early modETV2 (48 h) differentiation protocols. (**D**) Quantification on the percentage of h-iECs (CD31+) by flow cytometry for both differentiation protocols and each inhibitor.

**Fig. S7.**
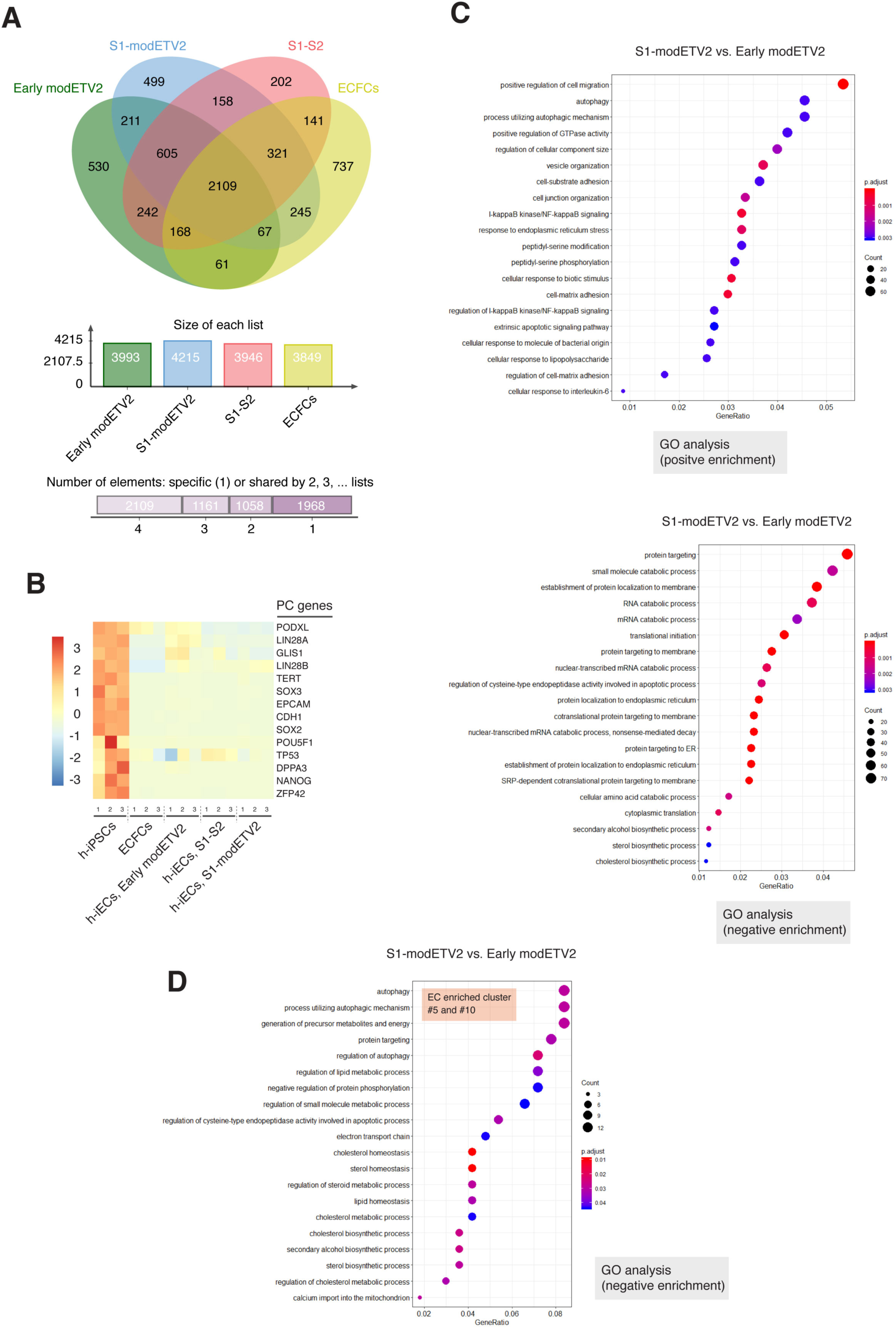
Transcriptome analysis of h-iECs obtained from various differentiation protocols. **(A)** Venn diagram on the number of differentially expressed genes in h-iECs and ECFCs that were upregulated compared to h-iPSCs. **(B)** Heatmap of the normalized expression of selected pluripotent genes across all samples. **(C)** GO analysis on all differentially expressed genes between h-iECs generated with the S1-modETV2 and the early modETV2 differentiation protocols. GO terms displayed corresponding to positive and negative enrichment for h-iECs generated with the S1-modETV2 protocol. **(D)** GO analysis on differentially expressed genes from EC clusters #5 and #10 between h-iECs generated with the S1-modETV2 and the early modETV2 differentiation protocols. GO terms displayed corresponding to the negative enrichment for h-iECs generated with the S1-modETV2 protocol.

**Fig. S8.**
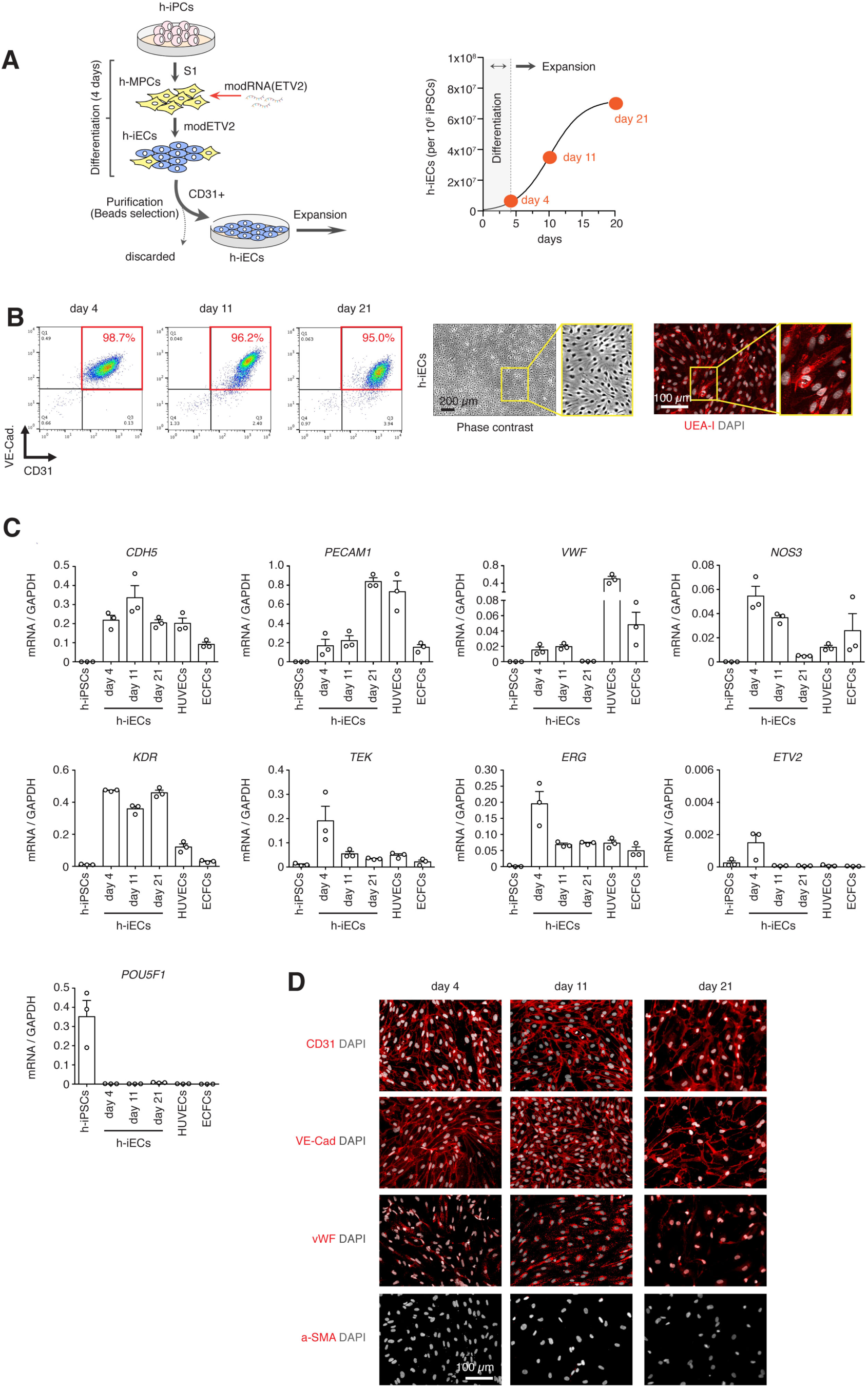
Robust endothelial phenotype of h-iECs along their expansion in culture. h-iECs generated by the S1-modETV2 protocol maintained an endothelial phenotype during *in vitro* expansion. (**A**) Schematic of S1-modETV2 protocol to generate, purify, and expand h-iECs. h-iECs were analyzed at three different time points of expansion (namely, day 4, day 11, and day 21). **(B)** Flow cytometry analysis revealed that h-iECs remained fairly pure during expansion (> 95% cells are VE-cadherin+/CD31+). Confluent h-iECs showed typical cobblestone morphology and UEA-I staining. (**C-D**) The expanded h-iECs maintained expression of EC markers at the mRNA **(C)** and protein (**D**) levels and remained negative for *POU5F1* (OCT4) and α-Smooth muscle actin (α-SMA).

**Fig. S9.**
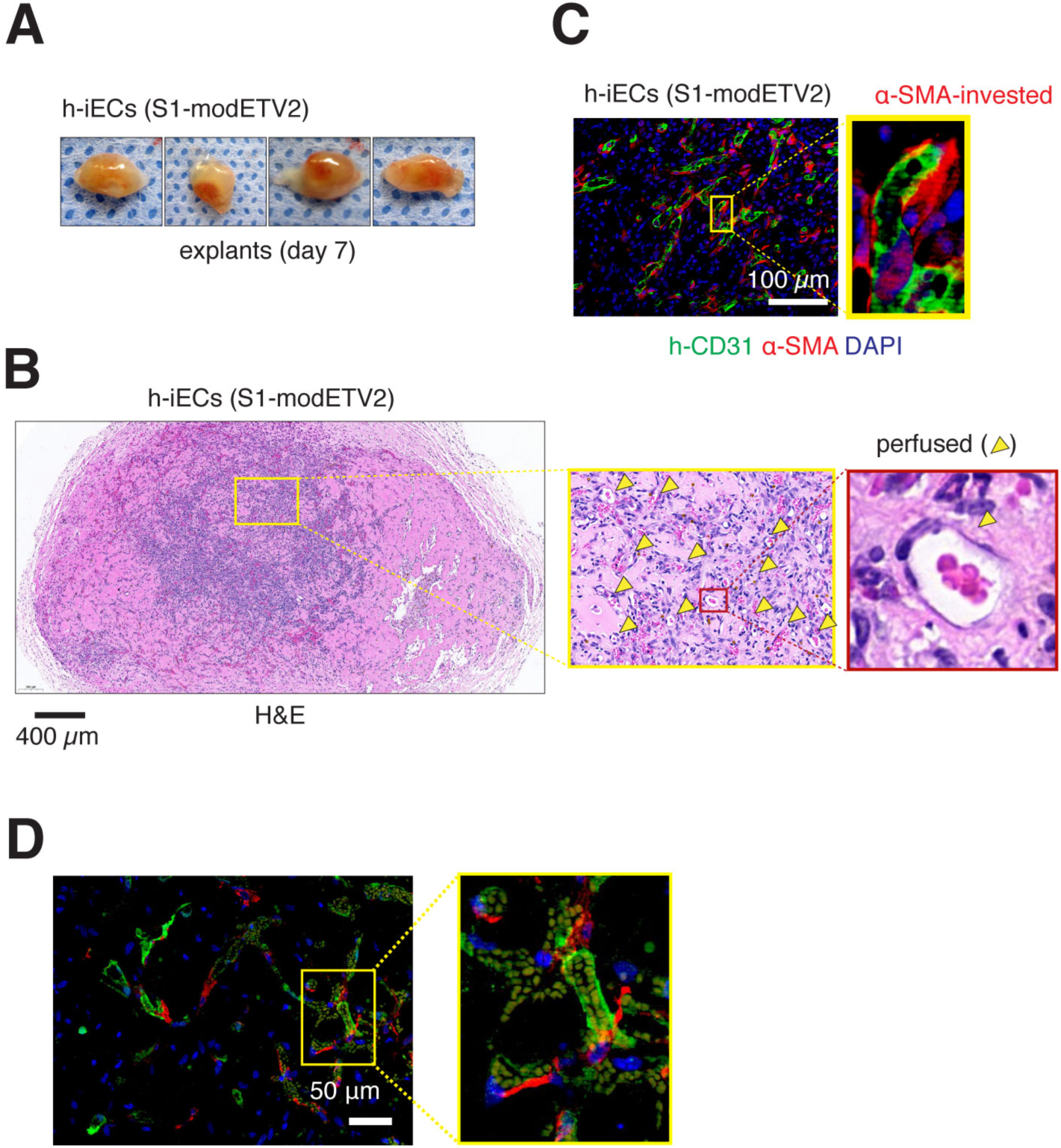
*In vivo* vascular network-forming ability of h-iECs without h-MSCs. (**A**) Grafts contained only h-iECs that were generated by the S1-modETV2 protocol and expanded until day 18. Images are macroscopic views of the explanted grafts from four mice at day 7. (**B**) Hematoxylin and eosin (H&E) staining of explanted grafts after 7 days *in vivo*. Perfused vessels were identified as luminal structures containing red blood cells (yellow arrowheads). Scale bar, 400 μm. (**C**) Immunofluorescent staining of explanted grafts after 7 days *in vivo*. Human lumens stained by h-CD31. Perivascular coverage stained by α-SMA. Nuclei stained by DAPI. Scale bar, 100 μm. (**D**) Immunofluorescence staining of explanted grafts that contained h-iECs-GFP after 7 days *in vivo*. Human lumens stained by GFP (green). Perivascular cells stained by α-SMA (red). Nuclei stained by DAPI. Scale bar, 50 μm.

**Table S1.**
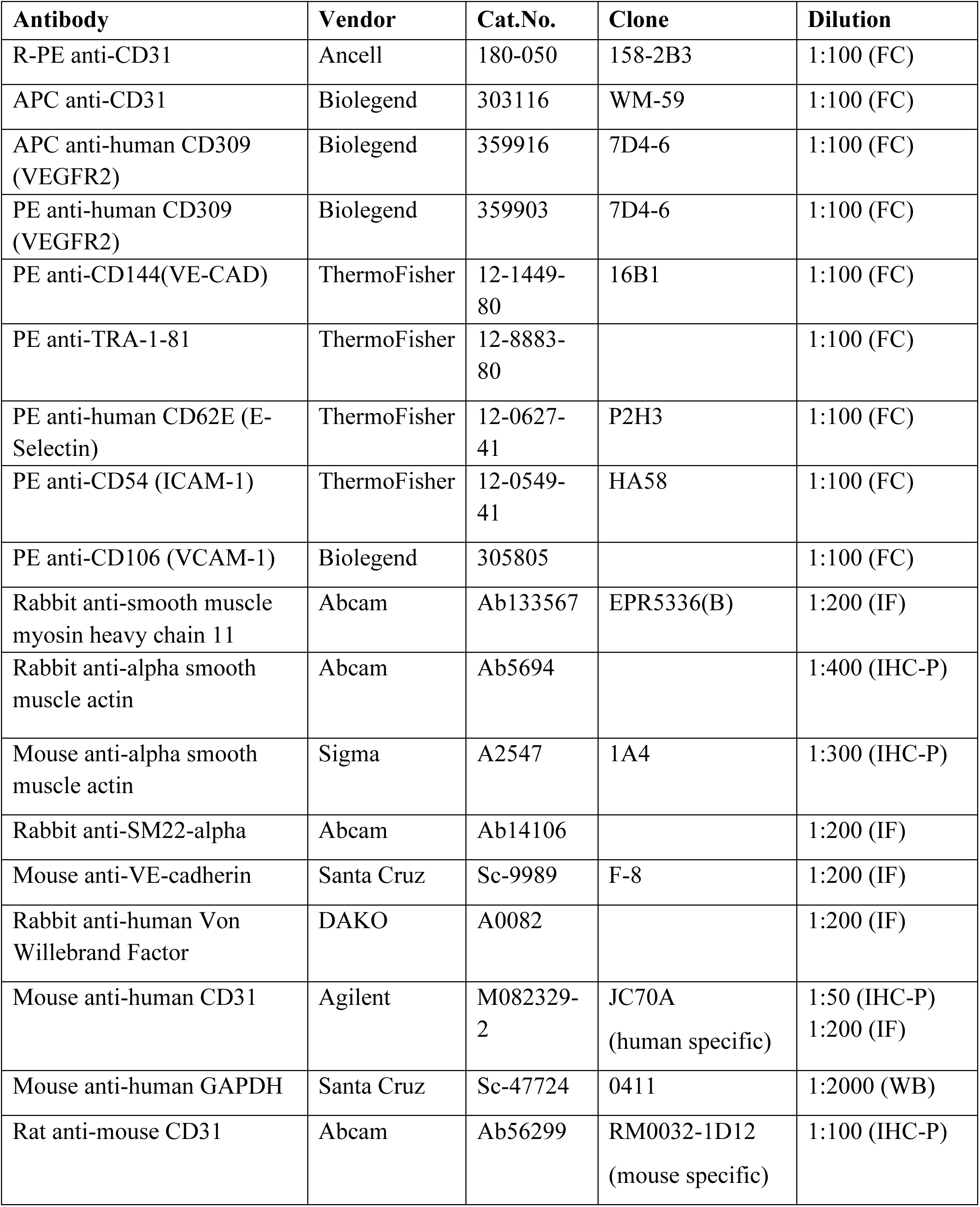
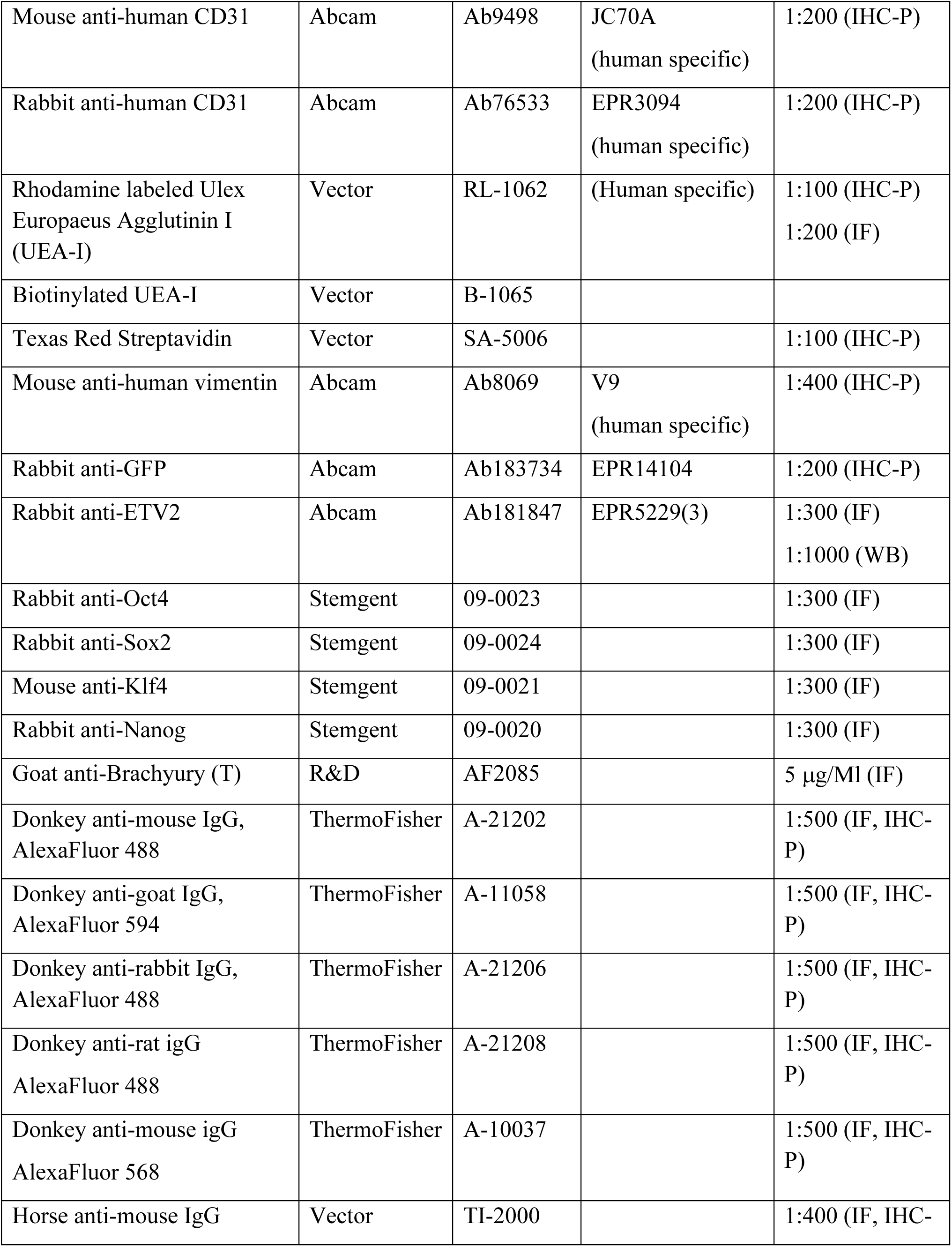
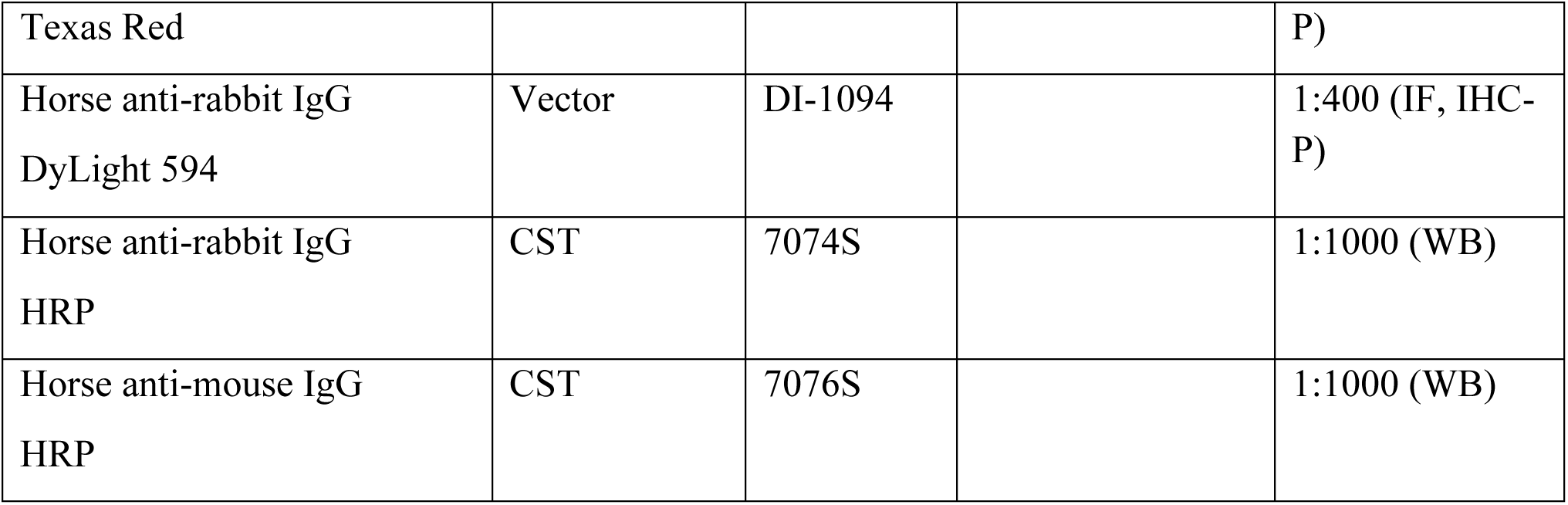
List of antibodies used in the study.

**Table S2.**
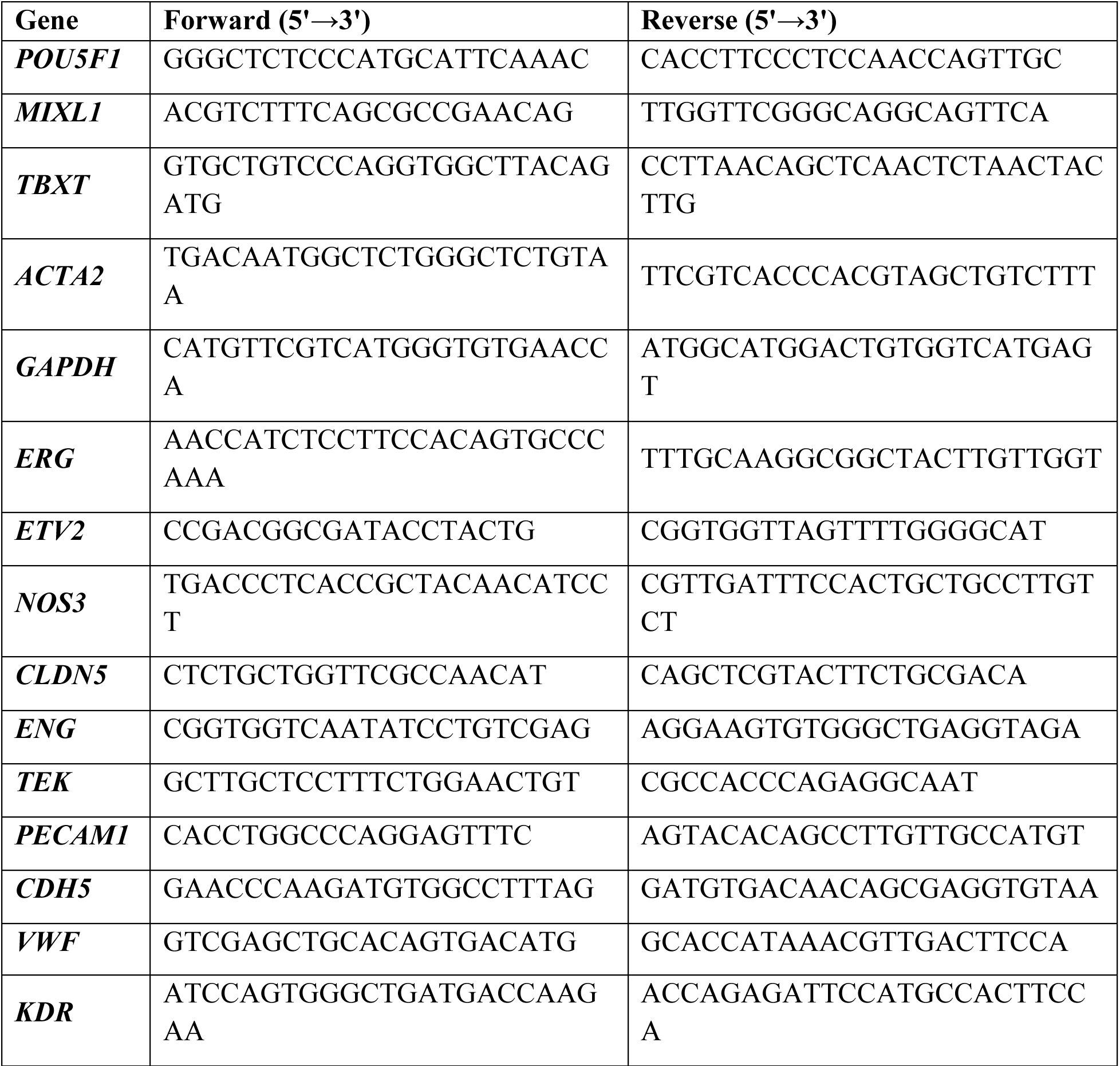
Sequences of primers used for qRT-PCR.

